# Cdh-2 and cortical f-actin dynamically cooperate to establish a stiffness gradient which contributes to forebrain roof plate invagination

**DOI:** 10.1101/2025.11.03.686203

**Authors:** Meenu Sachdeva, Prasenjit Sharma, Pankaj Gupta, Sweta Kushwaha, Mohd Ali Abbas Zaidi, Jonaki Sen

**Affiliations:** Department of Biological Sciences and Bioengineering, Indian Institute of Technology Kanpur, Kanpur, 208016, Uttar Pradesh, India; Mehta Family Center for Engineering in Medicine (MFCEM), Indian Institute of Technology Kanpur, Kanpur, 208016, Uttar Pradesh, India; Department of Mechanical Engineering, Indian Institute of Technology Kanpur, Kanpur, 208016, Uttar Pradesh, India; Department of Biochemistry and Molecular Biology, University of Nebraska Medical Center, Omaha, NE, 68198, USA

**Keywords:** Chick forebrain roof plate, Neuroepithelium, Overlying mesenchyme, Invagination, Stiffness

## Abstract

The invagination of the dorsal forebrain roof plate is critical for cerebral hemisphere formation. Although mutations in several genes have been linked to holoprosencephaly (HPE), and studies in chick embryos have implicated key signaling pathways, the mechanisms by which mechanical forces drive roof plate invagination remain poorly understood. To address this gap, we employed atomic force microscopy to map the spatiotemporal changes in tissue stiffness across the dorsal forebrain in the chick embryo during roof plate invagination. Our analysis revealed that neuroepithelium that is uniformly stiff initially, undergoes dramatic remodelling during this period of morphological change. Ultimately a pronounced stiffness gradient emerges, with the roof plate midline becoming markedly more compliant than the dorsolateral regions. Through expression screening, we identified Cdh2 as a candidate molecular regulator, whose spatiotemporal expression pattern closely mirrors the observed stiffness gradient. Mechanistically, we found that the interplay between Cdh2 and F-actin modulates tissue stiffness by regulating Cdh2 expression levels, apical adherens junction stability, and cortical F-actin distribution along the apico-basal axis of neuroepithelial cells. These findings underscore the importance of apico-basal localization of Cdh2 and provide critical insights into the mechanical forces and molecular interactions that govern forebrain roof plate morphogenesis, while also shedding light on the pathogenesis of HPE.

## Introduction

During embryonic development, the coordinated integration of signaling pathways and physical forces drives the precise formation of complex three-dimensional structures (K. E. Garcia et al., 2019; Iwashita et al., 2014; Liu et al., 2010). In vertebrates, following neural tube closure, the anterior neural tube expands to form three primary vesicles, with the most anterior vesicle giving rise to the forebrain. Subsequently, the dorsal region of the forebrain (roof plate) undergoes invagination and eventually meets the ventrally invaginating floor plate. This partitions the forebrain into two chambers that develop into the left and right cerebral hemispheres. Failure of this process leads to holoprosencephaly (HPE), a severe congenital brain malformation in humans that is quite common (Chafiq et al., 2024; Hamza & Higgins, 2017; Muenke, 2003). The dorsal forebrain consists of a pseudostratified neuroepithelium overlaid by a layer of neural crest derived mesenchyme that later contributes to meningeal formation. In the chick embryo, this bilayer structure undergoes the following morphological changes: A characteristic “pucker” first appears in the dorsal midline at Hamburger and Hamilton (HH) 19 (Hamburger & Hamilton, 1992) which subsequently bends inwards to form a distinctive W-shaped invagination by HH 23. Previous work from our laboratory demonstrated that the interplay between canonical and non-canonical BMP signaling establishes differential thickness of the roof plate neuroepithelium, a characteristic feature essential for the W-shaped invagination (Zaidi et al., 2025). We also showed that retinoic acid signaling regulates cell proliferation and patterning of the roof plate, which contribute to its invagination (Gupta & Sen, 2015). In addition, the extensive morphological changes occurring in the roof plate suggest that mechanical forces are likely to be involved in the invagination process. Recent advances in mechanobiology have highlighted that physical forces and molecular signals act in concert to shape embryonic tissues (K. Garcia et al., 2019). In particular, the mechanical properties such as tissue stiffness in the developing brain, have emerged as key regulators of morphogenesis. In fact, atomic force microscopy (AFM) based studies in the mouse cortex have revealed spatially and temporally regulated stiffness gradients that guide cellular behavior (Iwashita et al., 2014).

Nonetheless, several critical questions remain unresolved regarding chick forebrain roof plate invagination. For example, it remains to be determined whether mechanical forces contribute to this process and, if so, how this is integrated with molecular signals to control the precise timing and location of the invagination. Accumulating evidence indicates that the differential thickness of the neuroepithelium from midline to lateral regions arises, at least in part, from dynamic remodeling of the actin cytoskeleton (Zaidi et al., 2025). In addition to differential thickness, the forebrain roof plate also exhibits differential cell proliferation with the midline showing near absence of proliferating cells in contrast to the highly proliferating lateral regions (Gupta & Sen, 2015; Udaykumar et al., 2023). Thus, we hypothesize that the non-proliferative roof plate experiences tangential compression from the surrounding, highly proliferative neuroepithelium, possibly driving inward folding leading to invagination. These observations strongly implicate mechanical forces regulating this process.

In this study, we have investigated tissue stiffness and its role in translating molecular interactions into tissue-scale mechanical properties. Recent findings indicate that the relationship between cell adhesion and tissue mechanics is quite complex, as this involves dynamic interplay among adhesion molecules, cytoskeletal networks, and mechanical forces. In the context of the forebrain roof plate, the potential contributions of differential tissue stiffness and/or adhesion to invagination have never been explored. Here, we report that the classical cadherin N-cadherin (Cdh2) plays a critical role in regulating tissue stiffness and morphogenesis of the forebrain roof plate by modulating adherens junction stability and the cortical distribution of F-actin.

## Results

### Spatiotemporal changes in stiffness of the forebrain roof plate and overlying mesenchyme during morphogenesis

Visible morphological changes in the dorsal forebrain of the chick begins at embryonic day 3 (stage HH18). To observe these changes more closely, we harvested chick embryos at various stages and prepared coronal sections spanning the mid-anterior to mid-posterior regions of the forebrain (Figure 1A). These sections were stained with DAPI to label the nuclei, and we observed that the forebrain was elliptical in shape at HH18 (Fig. 1B and B’). However, at HH19, a “pucker” appeared in the middle of the neuroepithelium (NE) in the dorsal region known as the roof plate (Fig. 1C and C’) and the roof plate midline began to invaginate (move inwards) in the form of a shallow “W” by HH21 (Fig. 1D and D’). This invagination deepened further and at HH23 it was in the shape of a completely formed “W” (Fig. 1E and E’). In contrast, the morphology of the overlying mesenchyme remained relatively unchanged throughout this period, although there was a consistent increase in thickness from HH19 to HH23, particularly in regions just above the troughs of the W-shaped roof plate invagination, which we refer to as midline vortices (yellow asterisk in Fig. 1E’).

**Fig. 1.**
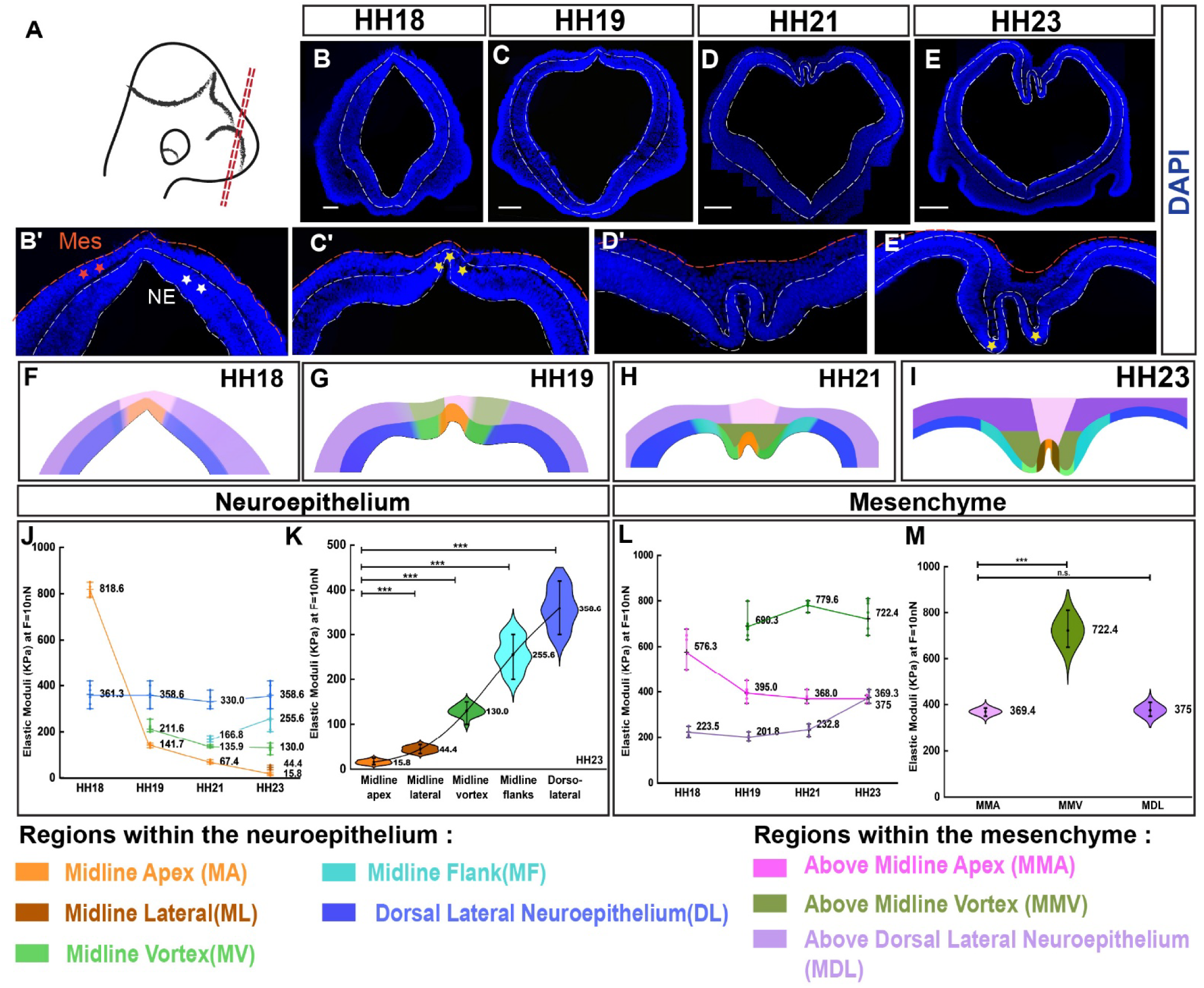
Spatiotemporal changes in tissue stiffness of the neuroepithelium and mesenchyme during dorsal forebrain morphogenesis. (A) A schematic illustration of the head of a chick embryo with the dashed red lines indicated the plane of sectioning of the forebrain. Coronal sections were taken from the mid-anterior to mid-posterior regions of the chick forebrain. (B,C,D,E) Images of DAPI-stained coronal sections from the mid-posterior region of the chick forebrain at various developmental stages from HH18 to HH23, 20X magnification; the white dashed line demarcates the neuroepithelium. Scale bar: 100µm (B′, C′, D′, E′) High magnification images of the DAPI stained dorsal forebrain sections at different stages of development from HH18 to HH23. The neuroepithelium is denoted by the white dashed line and asterisks, while the mesenchyme is represented by the red line and red asterisks. Yellow asterisks mark the “pucker” at HH19 (C’) and midline vortex at HH23 (E’). (F, G, H, I) Schematic of the dorsal forebrain at stages HH18, HH19, HH21 and HH23 with distinct colours marking subdomains of the roof plate neuroepithelium and overlying mesenchyme. The colour code for each subdomain is provided at the bottom. (J) The change in stiffness of different subdomains of the neuroepithelium plotted across developmental time. Each coloured line represents a distinct subdomain of the neuroepithelium. (N=5) (K) Violin plots of the stiffness of each subdomain of the neuroepithelium at HH23 (colour coded in I), with mean values and SEM indicated (N=8). The stiffness of the midline apex (MA) significantly different from other subdomains (p-value ≤ 0.001). (L) The change in stiffness of different subdomains of the dorsal mesenchyme plotted across developmental time. Each coloured line represents a distinct subdomain of the mesenchyme. (N=5) (M) Violin plots of the stiffness of each subdomain of the dorsal mesenchyme at HH23 (colour coded in I), with mean values and SEM indicated (N=8). The stiffness of the mesenchyme above midline apex (MMA) is significantly different from that above the midline vortex (MMV) (p-value ≤ 0.001).Two-way ANOVA was used for statistical analysis, followed by Tukey’s post-hoc analysis (K,M), with the p-values indicated. See also Fig. S1.

The morphological changes in the dorsal forebrain during invagination led us to hypothesize that these changes could be linked to concurrent changes in tissue stiffness. To establish if this is true, we proposed to independently quantify the stiffness of the dorsal forebrain neuroepithelium and the overlying mesenchyme between the developmental time windows of HH18 and HH23. In the absence of an established protocol for preparing chick (Gallus gallus) samples for atomic force microscopy (AFM), we first optimized the protocol for conducting contact-mode AFM on the embryonic chick forebrain (Fig. S1). The dorsal forebrain midline spans a substantial length of up to 1000 µm, while the contact area of the AFM probe covers only approximately 10 nm; hence, we divided the neuroepithelium and the overlying mesenchyme into distinct zones for systematic measurement (Fig. S1, S2, and S3). At HH18, the midline apex (MA) (Fig. 1F) of the neuroepithelium exhibited stiffness averaging at 818 kPa (Fig. 1J), while in the dorsolateral regions (DL), it was 361 kPa. On the other hand, the mesenchyme stiffness averaging at ∼ 576 kPa at the midline (MMA), although lower than that of the neuroepithelium at MA, it is quite elevated compared to the stiffness of the dorsolateral mesenchyme (MDL) averaging at 223.5kPa (Fig. 1L).

At HH19 (∼6 Hrs. later), the roof plate domain expands, and the “pucker” appears in the middle of the neuroepithelium (Fig. 1C’ and G). The midline apex (MA) of the neuroepithelium at HH19 exhibited a stiffness of ∼ 141 kPa, which was significantly lower than that of the MA region at HH18, whereas in the dorsolateral (DL) region, the stiffness remained nearly the same as that of HH18 (Fig. 1J). At this stage, the stiffness of the mesenchyme above the midline apex (MMA) decreased to 395 kPa; however, in the regions adjacent to the midline vortex (MMV), it was significantly elevated to nearly 690 kPa (Fig. 1L). Thus, the stiffness of the neuroepithelium exhibits a sharply decreasing trend, making it more flexible, while the stiffness of the overlying mesenchyme rises at sites where the neuroepithelium will undergo bending in future (Fig. 1G, J, and L). At HH21, the stiffness of the neuroepithelium at the midline apex (MA) further declined to 67 kPa, whereas at the dorsolateral (DL) regions, it remained largely unchanged, thus exhibiting a pronounced gradient of stiffness across the W-shaped region (Fig. 1J). In the mesenchyme, the areas directly above the midline vortices (MMV), where bends in the neuroepithelium are observed, exhibit significantly elevated levels of stiffness (∼780 kPa). In contrast, the stiffness of the mesenchyme in other regions, such as just above the midline apex (MMA) and above the dorsolateral neuroepithelium (MDL), remained at significantly lower levels (Fig. 1L). This gradient of stiffness across the invaginating neuroepithelium was quite apparent at HH23 (Fig. 1K), while the stiffness of the mesenchyme above the midline vortex (MMV) remained significantly elevated at ∼700 kPa at this stage (Fig. 1M).

The variations in tissue stiffness across the dorsal forebrain at different stages highlights the spatiotemporal dynamics of the mechanical properties of the neuroepithelium and overlying mesenchyme. We observed that with progress in development, the stiffness of the neuroepithelium, which was high in the midline and relatively low in the dorsolateral regions at HH18, exhibited an inverse gradient across the roof plate at HH23, such that the midline became considerably less stiff compared to the dorsolateral regions. The mesenchyme on the other hand, maintains moderate levels of stiffness above the dorsal neuroepithelium, except for the regions above the midline vortices (MMV) which become extremely stiff as the invagination of the neuroepithelium progresses. Thus, it is likely that the localized increase in stiffness of the mesenchyme provides the necessary force to drive invagination of the relatively more flexible neuroepithelium underneath.

### Cdh-2 exhibits a spatiotemporal expression pattern that aligns with changes in roof plate stiffness during morphogenesis

Spatiotemporal profiling of stiffness in the dorsal forebrain led us to speculate that roof plate invagination might be significantly influenced by mechanical forces that operate in real time along with biochemical pathways. In general, sites where cells adhere to each other serve as force-sensing hubs These sites are enriched in cadherins, which facilitate physical connections between neighboring cells through both trans- and cis-extracellular interactions and cytoplasmic associations with the actin cytoskeleton (Hu et al., 2019; Meng & Takeichi, 2009). We performed an expression screen using RNA *in situ* hybridization to identify candidate cell adhesion molecules involved in this process. We identified N-Cadherin/Cadherin-2 (Cdh-2) as a strong candidate, as it exhibited relatively high levels of expression in the W-shaped invaginating region of the roof plate compared to the dorsolateral regions of the neuroepithelium (Fig. 2A and A’). In the chick embryo, the walls of the forebrain anlagen are of variable thickness, comprising a single layer of neuroepithelial cells, with each cell spanning the apical-basal axis. The nuclei of neuroepithelial cells are positioned at different levels, giving it a multilayered appearance; hence, it is referred to as pseudostratified neuroepithelium. The apical side, adjacent to the lumen, contains apical adherens junctions, while the basal side is adjacent to the mesenchyme (Fig. 2B, B’ and B’’). We carried out immunohistochemistry on sections of the chick forebrain from stages spanning HH10 to HH23 to determine the expression of Cdh-2 and we found that between HH14 to HH18, Cdh-2 is specifically localized to the apical side of the neuroepithelium (Fig. 2C, D and E). From HH18 onwards, we observed that, in addition to being present in the apical side of all neuroepithelial cells, Cdh-2 protein could also be detected on the lateral and basal sides of those neuroepithelial cells present in the middle region of the roof plate (Fig. 2E, E’, F’, G, H, and H’). The distribution of Cdh-2 protein is shown schematically in Fig. 2J. The intensity of immunostaining progressively increased from HH18 to HH23 (Fig. 2I), indicating an increase in protein expression. It must be noted that although Cdh-2 exhibits differential expression across the roof plate neuroepithelium, we could not detect any expression of Cdh-2 in the mesenchyme. Interestingly, a strong correlation (R^2^ = 0.76) was observed when comparing the spatiotemporal changes in stiffness with the presence of the Cdh-2 protein in the midline apex region of the neuroepithelium (Fig. 2K). In fact, the prominent appearance of Cdh-2 protein at HH19 in the basal and lateral regions of cells in the neuroepithelium coincided with the dramatic decrease in stiffness of the neuroepithelium in this region (Fig. 2I, K, and L). These intriguing observations motivated us to further investigate the possible role of Cdh-2 in determining the stiffness of the roof plate and thereby regulating its invagination.

**Fig. 2.**
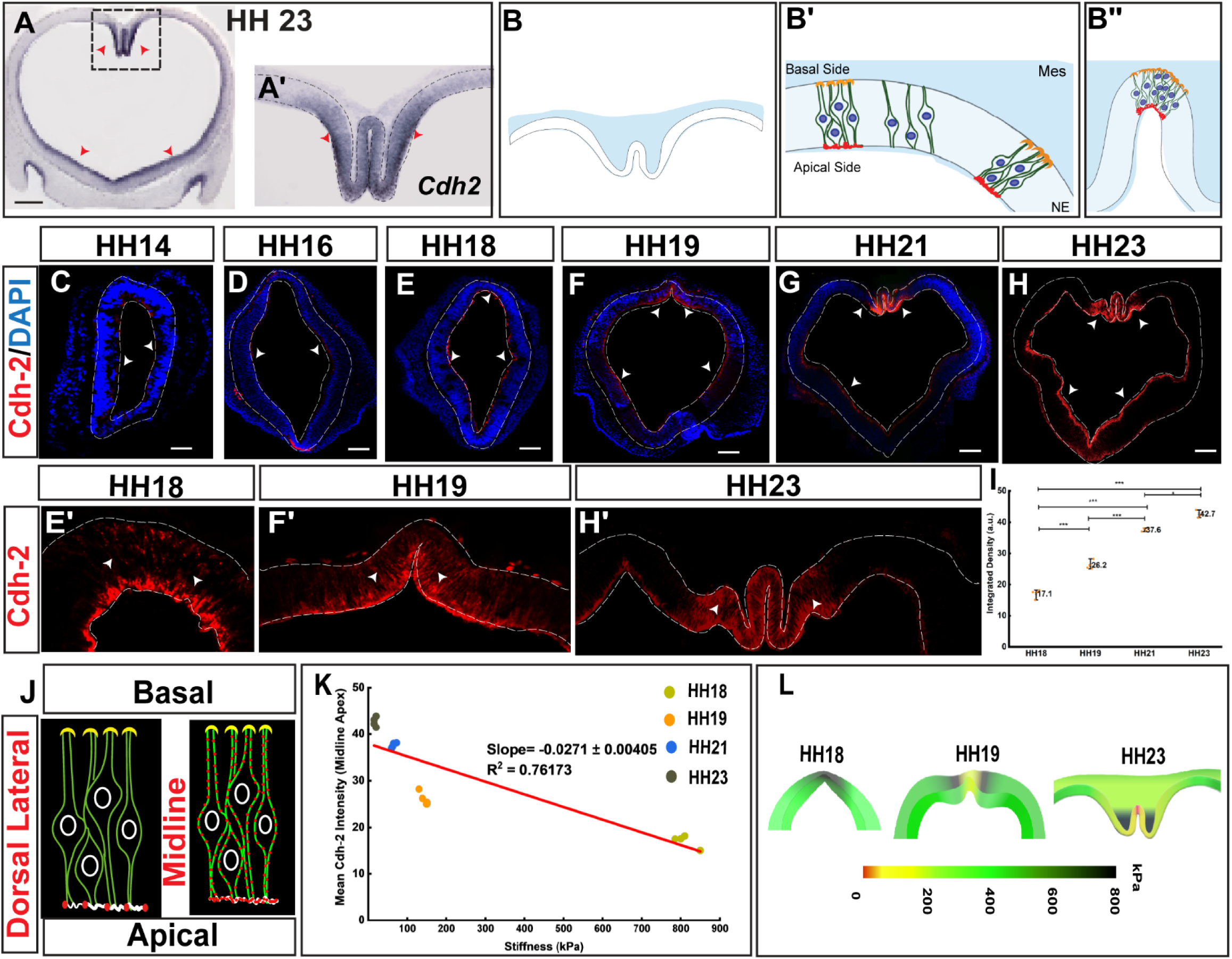
The dynamics of Cdh-2 expression correlate with stiffness changes during forebrain morphogenesis. (A) Sections of the chick forebrain at HH23 showing mRNA expression of *Cdh2.* The red arrowheads indicate domains where *Cdh2* expression is relatively high. Scale bar: 100 μm. (A′) High magnification image of the forebrain midline region at HH23 shown in the boxed region of (A). (B) A schematic representation of the dorsal forebrain neuroepithelium, the apical (red circles) and basal (orange circles) sides are indicated. (B’) A schematic representation of the pseudostratified neuroepithelium with every cell spanning the apical to the basal side and nuclei located at distinct depths. (B’’) A schematic representation of the pseudostratified neuroepithelium depicting the decrease in thickness from the dorsal lateral region to the midline of the roof plate. (C,D,E,F,H) Immunohistochemical detection of Cdh-2 in sections of the forebrain at different stages from HH14 (embryonic day 2) to HH23 (embryonic day 3.5). (C, D, E, F) DAPI (blue) and Cdh-2 (red) merged image of the section of chick forebrain from HH14 to HH21. (H) Image of a section of the chick forebrain at HH23 with Cdh-2 (red) immunostaining. A white arrow marks the region of high expression. Scale bar: 100 μm. (E’, F’, H’) High magnification images of sections of the dorsal forebrain with Cdh-2 immunostaining (red) at HH18, HH19, and HH23.\ (I) The change in the intensity of Cdh-2 staining in the roof plate midline plotted across developmental time (HH18 to HH23) (N=4). Significance of difference in Cdh-2 staining between stages assessed by unpaired t-test using Origin 2024b (p ≤ 0.001). (J) Schematic representation of distribution of Cdh-2 protein across the apicobasal axis of neuroepithelial cells in the roof plate midline and dorsal-lateral regions of the forebrain. The neuroepithelial cell membrane is coloured green and Cdh-2 protein is denoted by red circles. (K) Regression analysis curve depicting correlation between the change in Cdh-2 intensity and the change in stiffness over time at the midline apex region using Origin 2024b. (N=4) (L) Schematic illustration depicting the variation in stiffness (using a colour code) across different subdomains of the neuroepithelium and mesenchyme of the dorsal forebrain, during the invagination process at embryonic stages HH18, HH19, and HH23.

### Cdh-2 regulates invagination as well as patterning of the forebrain roof plate

We utilized loss-of-function (LOF) and gain-of-function (GOF) strategies to manipulate the function of Cdh-2 and gain insight into its possible role in the morphogenesis of the forebrain roof plate. For LOF, we generated a construct that expresses a dominant-negative version of Cdh-2 lacking the extracellular binding domain (DN-Cdh-2) (Kintner, 1992) and cloned it into the pCAG-IRES-GFP vector (Fig. S4). When expressed in excess, DN-Cdh-2 interferes with the interactions between the extracellular domains of adjacent cis- or trans-configured Cdh-2 molecules. In addition, we generated a construct for RNA interference-mediated knockdown of Cdh-2 (Cdh-2 RNAi) and cloned it into the pRmiR vector (Smith et al., 2009)(Fig. S4). The knockdown efficacy of Cdh-2 RNAi was determined using a sensor assay (Fig. S5). Upon electroporation of either DN-Cdh2 or Cdh-2 RNAi to create the LOF of Cdh-2 (Fig. 3A), we observed a dramatic morphological change in the roof plate, whereby it evaginated (protruded outwards) instead of undergoing invagination (bending inward) (compare Fig. 3B with 3C and Fig. 3D with 3E, respectively). This evagination phenotype was more pronounced in the case of the LOF generated through RNAi-mediated knockdown of Cdh-2 (Fig. 3C) than in the LOF generated through expression of DN-Cdh-2 (Fig. 3E). For gain-of-function (GOF), we co-electroporated a construct with full-length Cdh-2 (Cdh-2 FL) (Fig. S4) and a construct expressing GFP (pCAG-GFP) to mark electroporated cells in the chick forebrain. Electroporation was conducted at HH10, and the tissue was harvested at HH23 (Fig. 3A). We observed that upon gain of function of Cdh-2, the morphology of the forebrain roof plate was significantly perturbed, and the invagination was V-shaped, as opposed to the W-shaped invagination in the control (Fig. 3F and G).

**Fig. 3.**
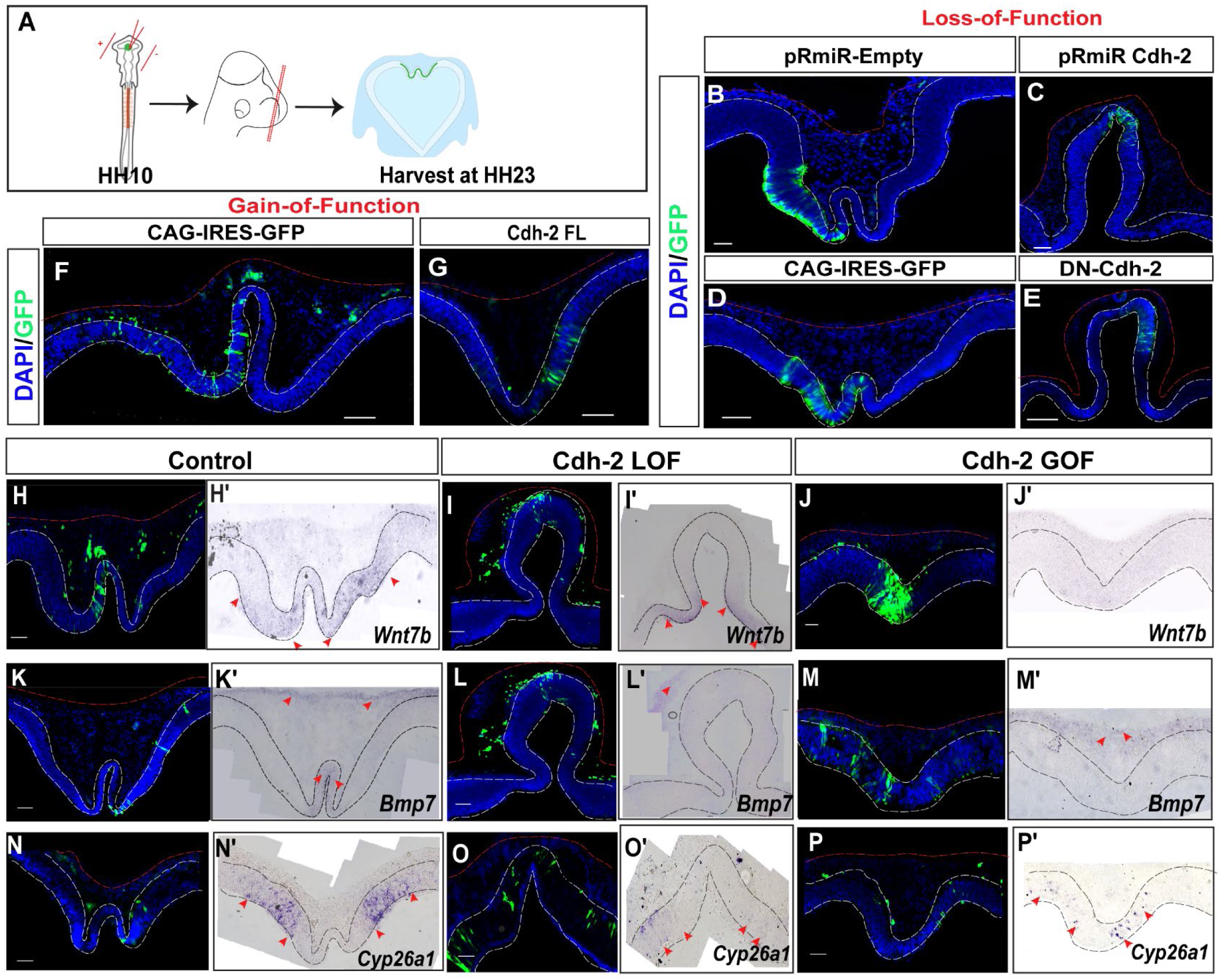
Disruption of Cdh-2 function impairs patterning of the roof plate. (A) Schematic of experimental design for generating LOF of Cdh-2 through electroporation of construct expressing dnCdh-2 or RNAi targeting Cdh-2 and GOF of Cdh-2 through electroporation of construct expressing Cdh-2 FL. (B,C) Images of DAPI stained sections of the forebrain roof plate of the chick embryo electroporated with control construct, pRmiR empty (B) and pRmiR-Cdh-2 RNAi (C) for LOF of Cdh-2. Green fluorescence marks the domain of electroporation. Scale bar: 100 μm. (D,E) Images of DAPI stained sections of the forebrain roof plate of the chick embryo electroporated with control construct, CAG-IRES-GFP (D) and pCAG-DN-Cdh2 (E) for LOF of Cdh-2. Green fluorescence marks the domain of electroporation. Scale bar: 100 μm. (F,G) Images of DAPI stained sections of the forebrain roof plate of the chick embryo electroporated with control construct, CAG-IRES-GFP (F) and pCAG- Cdh2-FL (G) for GOF of Cdh-2. Green fluorescence marks the domain of electroporation. Scale bar: 100 μm. (H,K,N) Images of DAPI stained sections of the chick forebrain roof plate at HH23 electroporated with the control construct pCAG-GFP. Green fluorescence indicates the extent of electroporation. (H’, K’, N’) Images of the chick forebrain roof plate at HH23 electroporated with the control construct pCAG-GFP with expression of *Wnt7b* (H’), *Bmp7* (K’), and *Cyp26a1*(N’) mRNA indicated by the purple-blue colour. Red arrows demarcate the domains of expression. Scale bar: 100 μm (N=3). (I, L, O) Images of DAPI stained sections of the chick forebrain roof plate at HH23 electroporated with the construct pCAG-DN-Cdh-2. Green fluorescence indicates the extent of electroporation. (I’, L’, O’) Images of the chick forebrain roof plate at HH23 electroporated with the construct pCAG-DN-Cdh-2 showing expression of *Wnt7b* (I’), *Bmp7* (L’), and *Cyp26a1*(O’) mRNA indicated by the purple-blue colour. Red arrows demarcate the domains of expression. Scale bar: 100 μm (N=3). (J,M,P) Images of DAPI stained sections of the chick forebrain roof plate at HH23 electroporated with the construct pCAG-Cdh-2-FL. Green fluorescence indicates the extent of electroporation. (J’, M’, P’) Images of the chick forebrain roof plate at HH23 electroporated with the construct p-CAG-Cdh-2-FL showing expression of *Wnt7b* (J’), *Bmp 7* (M’), and *Cyp26a1* (P’) mRNA indicated by the purple-blue colour. Red arrows demarcate the domains of expression. Scale bar: 100 μm (N=3).

In a previous study carried out by our group, we observed dramatic changes in forebrain roof plate invagination upon manipulation of RA signaling. These morphological changes were accompanied by alterations in the patterning of the roof plate, as gauged by differences in the expression patterns of certain roof plate marker genes (Gupta & Sen, 2015). Thus, we were curious to determine whether the LOF and/or GOF of Cdh-2 would affect patterning of the forebrain roof plate. We examined the expression of various markers such as WNT7B, BMP7, and CYP26A1 and found that the expression of WNT7B was completely lost with the GOF of Cdh-2 (Fig. 3H, H’, J, and J’), whereas with LOF it was present in the lateral regions of the evaginated roof plate (Fig. 3H, H’, I, and I’). BMP7, which is expressed only in the middle loop of the W-shaped invagination in the control, was completely absent in both the LOF and GOF of Cdh-2 (Fig. 3K, K’, L, L’, M, and M’). Further, with both LOF and GOF of Cdh-2 we observed a dramatic reduction in the expression of CYP26A1, which became restricted to very few cells (Fig. 3N, N’, O, O’, P, and P’). Overall, we observed a significant change in the marker expression and patterning of the dorsal forebrain upon functional perturbation of the Cdh-2.

### Cdh-2 mediates spatiotemporal variability in stiffness of the roof plate during its invagination

We observed that spatiotemporal changes in the stiffness of the dorsal forebrain tissues were closely correlated with the expression of Cdh-2 in the neuroepithelium. Further, upon perturbation of Cdh-2, morphology was severely perturbed. Together, these findings prompted us to investigate the effect of Cdh-2 on tissue stiffness. We measured stiffness using AFM after electroporation with Cdh-2 loss-of-function (LOF) and gain-of-function (GOF) constructs at HH23 (Fig. S6) when the W-shaped invagination was prominent in the dorsal forebrain of the chick embryo (Fig. 4A and A’). In the LOF condition at HH23, the stiffness was uniformly elevated across the evaginated roof plate neuroepithelium, averaging approximately 308 kPa at the midline apex and 358 kPa in the midline lateral regions. These values approached levels comparable to those of the dorsolateral neuroepithelium of the control roof plate at the same stage, which measured 389 kPa (Fig. 4B, B’, D, F, G, and H). This suggests that the LOF of Cdh-2 decreased the stiffness gradient normally present from the midline to the dorsolateral regions of the neuroepithelium across the dorsal forebrain (Fig. 4F, G, and H). Contrary to our expectations, GOF of Cdh-2 resulted in abnormally elevated midline stiffness relative to controls (Fig. 4C, C’, D, F, G, and H). In contrast, dorsolateral neuroepithelial stiffness was comparable to controls in both LOF and GOF conditions (Fig. 4D, F, G, and H).

**Fig. 4.**
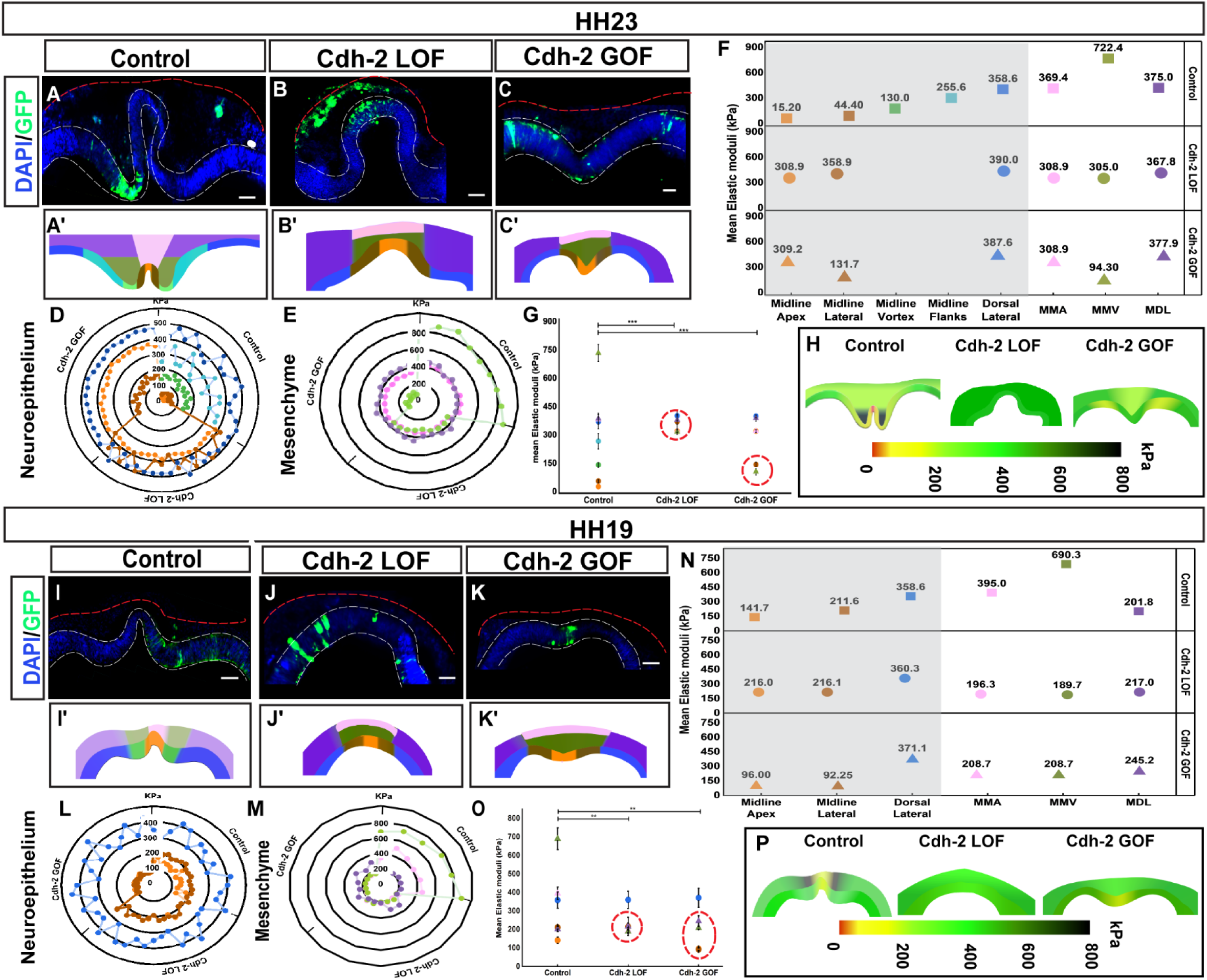
Modulation of Cdh-2 alters the stiffness of the neuroepithelium and overlying mesenchyme in the dorsal forebrain. (A,B,C) Images of DAPI stained sections of the chick forebrain roof plate at HH23 electroporated with control (pCAG-GFP), Cdh-2 LOF (pCAG-DN-Cdh-2) and Cdh-2 GOF (pCAG-Cdh-2-FL), respectively, the white dashed line demarcates the neuroepithelium Scale bar: 100 μm. (A’,B’,C’) A schematic representation of the chick forebrain roof plate at HH23 electroporated with control (A’), Cdh-2 LOF (B’) and Cdh-1 GOF (C’). The subdomains of the neuroepithelium and mesenchyme that were probed with AFM for measuring stiffness are denoted in different colours. (D) The radar plot comparing the stiffness of the roof plate neuroepithelium at HH23 in the control, Cdh-2 LOF and Cdh-2 GOF electroporated forebrains. Each dot represents the mean of the 100 points of region of interest (ROI) as shown in Fig. S6. (E) The radar plot comparing the stiffness of the dorsal mesenchyme at HH23 in the control, Cdh-2 LOF and Cdh-2 GOF electroporated forebrains. Each dot represents the mean of the 100 points of each region of interest (ROI) as shown in Fig. S6. (F) The scatter plot for the mean stiffness of each ROI of the roof plate neuroepithelium (grey shaded region) and the dorsal mesenchyme (unshaded region) at HH23 of control, Cdh-2 LOF, and Cdh-2 GOF electroporated forebrains. Each dot represents the mean of the 100 points of each region of interest (ROI) as shown in Fig. S6. (G) The dot plot for the mean stiffness (Mean ± SEM) of each ROI of the roof plate neuroepithelium (grey shaded region) and the dorsal mesenchyme (unshaded region) at HH23 of control, Cdh-2 LOF, and Cdh-2 GOF electroporated forebrains. Colour coded circles and triangles represent each ROI in the neuroepithelium and mesenchyme, respectively. Each circle/triangle represents the mean of the 100 points within each region of interest (ROI) as shown in Fig. S6. Red dashed circles indicate the distribution of the mean stiffness across the subdomains of the roof plate neuroepithelium and the dorsal mesenchyme. Unpaired t-test using Origin 2024b was used to determine significance p ≤ 0.001. (N=4). (H) Schematic depicting the variation in stiffness (using a colour code) across different subdomains of the neuroepithelium and mesenchyme of the dorsal forebrain, upon Cdh-2 perturbation, during the invagination process at HH23. (I, J, K) Images of DAPI stained sections of the chick forebrain roof plate at HH19 electroporated with control (pCAG-GFP), Cdh-2 LOF (pCAG-DN-Cdh-2) and Cdh-2 GOF (pCAG-Cdh-2-FL), respectively, the white dashed line demarcates the neuroepithelium Scale bar: 100 μm. (I’, J’, K’) A schematic representation of the chick forebrain roof plate at HH19 electroporated with control (I’), Cdh-2 LOF (J’) and Cdh-1 GOF (K’). The subdomains of the neuroepithelium and mesenchyme that were probed with AFM for measuring stiffness are denoted in different colours. (L) The radar plot comparing the stiffness of the roof plate neuroepithelium at HH19 in the control, Cdh-2 LOF and Cdh-2 GOF electroporated forebrains. Each dot represents the mean of the 100 points of region of interest (ROI) as shown in Fig. S7. (M) The radar plot comparing the stiffness of the dorsal mesenchyme at HH19 in the control, Cdh-2 LOF and Cdh-2 GOF electroporated forebrains. Each dot represents the mean of the 100 points of each region of interest (ROI) as shown in Fig. S7. (N) The scatter plot for the mean stiffness of each ROI of the roof plate neuroepithelium (grey shaded region) and the dorsal mesenchyme (unshaded region) at HH19 of control, Cdh-2 LOF, and Cdh-2 GOF electroporated forebrains. Each dot represents the mean of the 100 points of each region of interest (ROI) as shown in Fig. S7. (O) The dot plot for the mean stiffness (Mean ± SEM) of each ROI of the roof plate neuroepithelium (grey shaded region) and the dorsal mesenchyme (unshaded region) at HH19 of control, Cdh-2 LOF, and Cdh-2 GOF electroporated forebrains. Colour coded circles and triangles represent each ROI in the neuroepithelium and the mesenchyme, respectively. Each circle/triangle represents the mean of the 100 points within each region of interest (ROI) as shown in Fig. S7. Red dashed circles indicate the distribution of the mean stiffness of subdomains of the roof plate neuroepithelium and the dorsal mesenchyme. Unpaired t-test using Origin 2024b was used to determine significance p ≤ 0.001. (N = 4). (P) Schematic depicting the variation in stiffness (using a colour code) across different subdomains of the neuroepithelium and mesenchyme of the dorsal forebrain, upon Cdh-2 perturbation, during the invagination process at HH19.

Interestingly when assessing the stiffness of the mesenchyme following LOF and GOF of Cdh-2 cells, we observed a significant decline in stiffness compared to the control. It is important to note that Cdh-2 is expressed in the roof plate neuroepithelium alone and is completely absent from the mesenchyme. Normally, at HH23, mesenchymal stiffness is relatively uniform, except in the region above the midline vortices, where it is significantly high at approximately 700 kPa (Fig. 4E, F, G, and H). However, with the LOF of Cdh-2 in the neuroepithelium, the mesenchymal stiffness above the vortices decreased to approximately 305 kPa, which is close to that of the dorsolateral mesenchyme, resulting in overall reduced mesenchymal stiffness. On the other hand, with GOF of Cdh-2, stiffness of the mesenchyme in the vortex region overlying the midline was also dramatically reduced to approximately 94.3 kPa, possibly leading to the mesenchyme exerting much less force on the underlying neuroepithelium (Fig. 4E, F, and H). This raises the following question: Under the GOF condition, how was the neuroepithelium undergoing bending to give rise to the V-shaped invagination if the overlying mesenchyme was much less stiff and, hence, could not apply the necessary force?

Our earlier observations suggest that invagination of the forebrain roof plate is initiated by the formation of the “pucker” at HH19. Subsequently, the neuroepithelium on either side of this “pucker” bends inward to give rise to the W-shaped morphology of the invagination at HH23. Therefore, we speculated that the morphological consequences of the LOF and GOF of Cdh-2 observed at HH23 could result from the disruption of Cdh-2 function at HH19. To explore this, we analyzed morphological changes at HH19 after Cdh-2 perturbation. LOF resulted in a flattened neuroepithelium lacking any discernible pucker (Fig. 4I and J), whereas GOF induced a subtle midline dip that likely develops into a V-shaped invagination at later stages (Fig. 4I and K). We therefore wondered whether Cdh-2 modulation also affects tissue stiffness at this stage. We measured the stiffness of the neuroepithelium and mesenchyme at HH19, following the LOF and GOF of Cdh2 (Fig. S7). We found that, following LOF at HH19, stiffness across the dorsal neuroepithelium was uniform (∼216 kPa), unlike in the control, where the stiffness was graded with the midline apex being much less stiff (∼ 41 kPa) than the midline lateral region (∼ 211 kPa) (Fig. 4L, N, O, and P). Similarly, with the GOF of Cdh-2, the stiffness of the dorsal neuroepithelium was close to 96 kPa, which is lower than the stiffness at both the midline apex and midline lateral regions in the control (Fig. 4L, N, O, and P). This indicates that GOF of Cdh-2 causes both a decrease in overall stiffness and a complete loss of the stiffness gradient across the neuroepithelium. Moreover, consistent with our observations at HH23, disruption of Cdh-2 significantly altered the stiffness of the overlying mesenchyme at HH19, resulting in an overall reduction in both LOF and GOF conditions compared to controls. The strong correlation between Cdh-2 expression, morphological defects, and disrupted stiffness gradients at HH19 further supports the conclusion that Cdh-2 is essential for maintaining proper stiffness in the dorsal forebrain during morphogenesis.

Furthermore, we found that LOF of Cdh-2 leads to evagination at HH23, even though both the mesenchyme and neuroepithelium exhibit uniform stiffness. In contrast, GOF causes bending of the neuroepithelium, resulting in a V-shaped invagination at HH23, despite markedly reduced stiffness in the mesenchyme overlying the midline. Detailed quantitative analysis showed that LOF of Cdh-2 reduced neuroepithelial stiffness to ∼216 kPa and mesenchymal stiffness to ∼189 kPa (Fig. 4L, M, N, and O). Therefore, the mesenchyme was less stiff than the neuroepithelium at HH19, which could explain why it evaginated (Fig. 4P). Conversely, GOF resulted in a higher mesenchymal stiffness (∼208 kPa), providing the necessary force for neuroepithelial bending. Simultaneously, it lowered the stiffness of the midline neuroepithelial region to ∼92 kPa, making it less stiff than the dorsolateral region, thus leading to a V-shaped invagination (Fig. 4P). The morphological defects in the roof plate — evagination upon LOF and V-shaped invagination upon GOF — likely result from a failure to establish the stiffness gradient required for pucker formation at HH19. These abnormalities therefore appear to arise from disrupted or misdirected mechanical cues due to functional perturbation of Cdh-2. Overall, our findings identify Cdh-2 as a key molecular player responsible for establishing the stiffness gradients necessary for proper invagination, as its perturbation affects both morphology and tissue stiffness in both LOF and GOF conditions.

Although distinctive morphological phenotypes were observed under Cdh-2 LOF and GOF conditions, the overall changes in tissue stiffness were remarkably similar. Both the LOF and GOF abolished the stiffness gradient extending from the midline of the roof plate neuroepithelium, resulting in a more or less uniform stiffness profile. To investigate the mechanism by which LOF and GOF manipulations produce similar changes in tissue stiffness, we performed immunohistochemistry for Cdh-2 on control and GOF forebrain sections at HH23. Contrary to our expectation that GOF would increase Cdh-2 expression, we observed a significant decrease in Cdh-2 immunostaining in the electroporated regions of GOF embryos compared with controls. Quantification of Cdh-2 expression in GOF samples at HH23 confirmed significantly reduced levels (Fig. S8A, A’, B, B’, C). At HH19, forebrain sections electroporated with GOF constructs showed no detectable Cdh-2 expression (Fig. S8D, D’, E, E’, F, F’). Based on these observations, we hypothesize that a negative feedback mechanism is activated in response to Cdh-2 overexpression. This leads to downregulation of endogenous Cdh-2 at HH19, followed by only partial recovery at HH23 that remains insufficient to restore proper roof plate flexibility, similar to the effect observed in Cdh-2 loss-of-function (LOF), explaining the similar neuroepithelial stiffness changes observed under both LOF and GOF conditions. These findings can also be extended to the progressive increase in apicobasal Cdh-2 distribution along the cell surface observed between HH18 and HH23 (Fig. 2I). Notably, this increase in Cdh-2 coincides with the stiffness changes (Fig. 2K). However, it remains unclear how the progressive apicobasal accumulation of Cdh-2 contributes to increasing the flexibility of the dorsal forebrain.

### Cdh2 translates mechanical cues through adherens junctions and not through direct regulation of cell proliferation

Certain tissue contexts have a high cell density and greater connectivity, such that cells are more tightly packed and interconnected, which restricts their movement. This collective behavior leads to a solid-like state, in which the tissue has increased mechanical rigidity and resists deformation. On the other hand, a lower cell density or reduced connectivity results in a more fluid-like state, where the tissue flows and cells have greater motility (Méhes & Vicsek, 2014). Cell density can be regulated by factors such as cell proliferation and death; hence, differential proliferation rates within a tissue may induce tissue buckling, leading to invagination of the forebrain roof plate midline.

At HH23, the chick forebrain neuroepithelium is highly proliferative throughout, except for the dorsal forebrain midline, corresponding to the middle loop of the W-shaped invagination (Gupta & Sen, 2015; Udaykumar et al., 2023). This indicates that differential proliferation is a distinctive feature of the neuroepithelium of the dorsal forebrain. To determine whether Cdh-2 plays a role in the establishment of differential cell proliferation in this context, we perturbed its function and assessed its effect on cell proliferation by immunohistochemical detection of phospho-histone 3 (PH3), which marks the mitotic phase of the cell cycle (Fig. S9A, D, B, E, C, F). Quantification of PH3-positive cells in both the midline and dorsolateral neuroepithelium following LOF of Cdh-2 revealed a significant increase in proliferation at the midline compared with the control. However, gain-of-function (GOF) perturbation did not produce a significant change in the number of proliferating cells in either the midline or dorsolateral regions (Fig. S9G). As Cdh-2 perturbation also alters neuroepithelial morphology in the dorsal forebrain, we investigated whether these morphological defects were a secondary consequence of changes in proliferation. Therefore, we perturbed Cdh-2 in the lateral neuroepithelium, followed by the assessment of cell proliferation through immunohistochemical detection of PH3 (Fig. S9H, I, J, H). Quantification of GFP and PH3 double-positive cells during LOF and GOF in the lateral neuroepithelium revealed no change in proliferation (Fig. S9L) or morphology at this location. Thus, the observed effect on cell proliferation in the roof plate likely resulted from context-dependent morphological and patterning changes caused by functional perturbation of Cdh-2 in the roof plate.

To understand how Cdh-2 links molecular changes to the mechanical properties, we examined adherens junctions, where high levels of Cdh-2 are observed. ZO-1 localizes to cadherin-based contacts through direct interaction with the cadherin–catenin complex (Fig. 5A). It scaffolds cadherin adhesion to the actin cytoskeleton and modulates mechanical forces at apical adherens junctions (Fanning et al., 2012; Itoh et al., 1997). In the absence of ZO-1, cadherin-containing junctions form but fail to properly connect to the cytoskeleton, weakening mechanical forces and stability (Maiers et al., 2013). We observed that ZO-1 immunostaining closely localized with Cdh-2 throughout the apical side of the forebrain neuroepithelium (Fig. 5B, B’, C, C’, D, and D’). This led us to examine the effect of Cdh-2 perturbation on ZO-1 at the apical adherens junctions. In both LOF and GOF for Cdh-2, immunohistochemical detection of ZO-1 expression revealed decreased levels within the electroporated regions compared to the control (Fig. 5E, E’, G, and G’), with a more pronounced reduction in ZO-1 observed with GOF for Cdh-2 (Fig. 5E, E’, F, and F’). Thus, adherens junctions appear to be disrupted by either the LOF or the GOF of Cdh-2. ZO-1 is essential for maintaining Cdh-2 at apical adherens junctions and for preserving tissue shape. This is selectively impaired by the depletion of ZO-1, leading to destabilized mechanical properties consistent with the stiffness changes observed with both the LOF and GOF of Cdh-2. Taken together, these results indicate that Cdh-2 controls cell-cell adhesion dynamics and mechanical stability at apical junctions through ZO-1, which is essential for proper forebrain roof plate invagination.

**Fig. 5.**
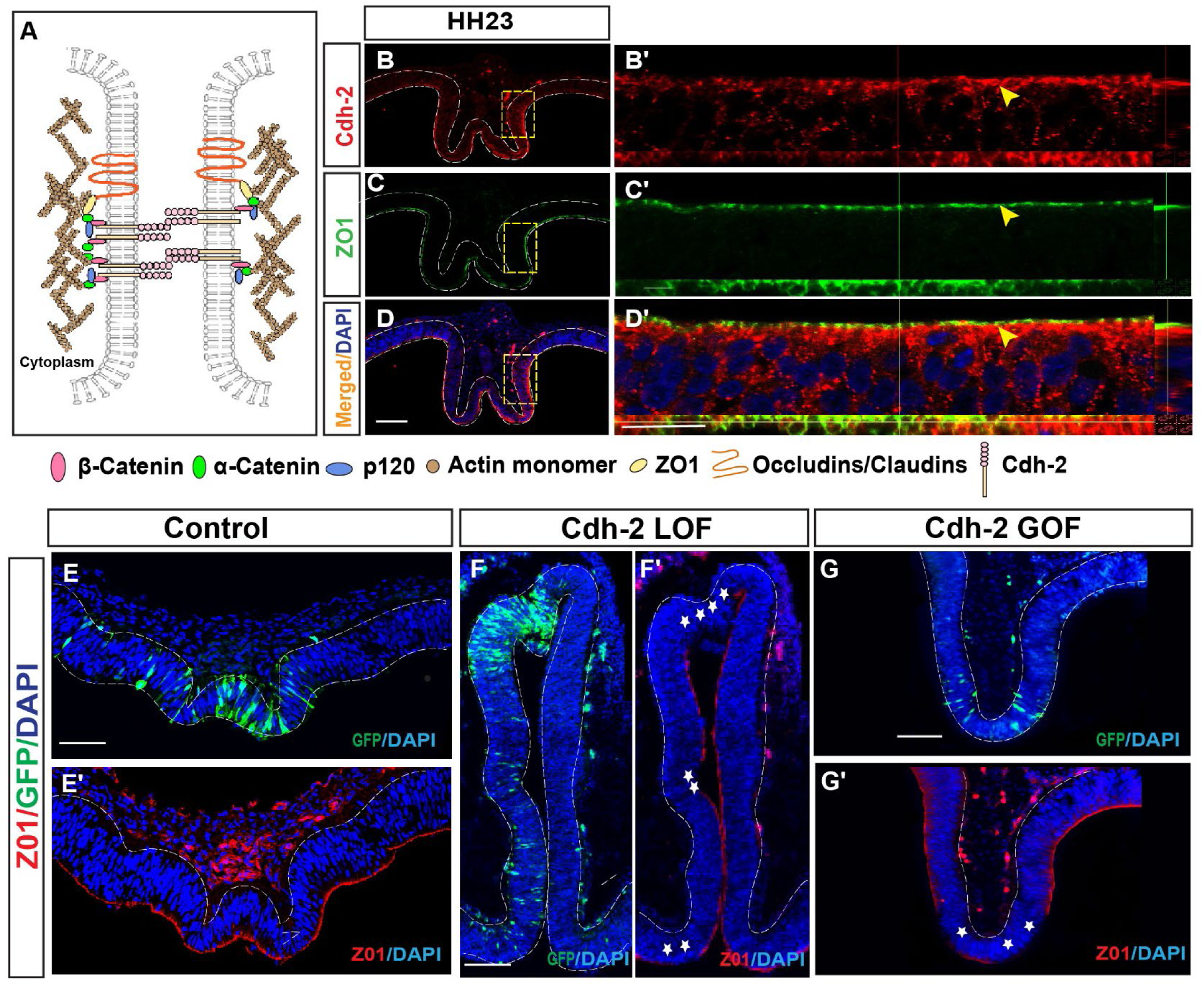
Functional manipulation of Cdh-2 disrupts the integrity of adherens junctions. (A) Schematic representation of adherence junction showing the ZO-1 and Cdh-2 and their interaction with the actin cytoskeleton. (B) Image of the section of the chick forebrain at HH23 with Cdh-2 immunohistochemistry (Red). White dashed line demarcates the neuroepithelium and yellow dashed box encloses the region of interest from which the high magnification (60X) image has been captured. Scale bar: 100µm. (B’) A frame from the images taken with a confocal microscope from the region within the yellow dashed box with Cdh-2 immunohistochemistry (red). Yellow arrow indicates the apical surface with Cdh-2 distribution. (C) Image of the section of the chick forebrain at HH23 with ZO-1 immunohistochemistry (green). White dashed line demarcates the neuroepithelium and yellow dashed box encloses the region of interest from which the high magnification (60X) image has been captured. Scale bar: 100µm (C’) A frame from the images taken with a confocal microscope from the region within the yellow dashed box with ZO-1 immunohistochemistry (green). Yellow arrow indicates the apical surface with ZO-1 distribution. (D) Merged Image of the section of the chick forebrain at HH23 with Cdh-2 (Red) and ZO-1(green) immunohistochemistry and DAPI stained nuclei (blue). White dashed line demarcates the neuroepithelium and yellow dashed box encloses the region of interest from which the high magnification (60X) image has been captured. Scale bar: 100µm (D’) A frame from the images taken with a confocal microscope from the region within the yellow dashed box with merged Cdh-2 (red) and ZO-1 (green) immunohistochemistry and DAPI stained nuclei (blue). Yellow arrow indicates the apical surface with Cdh-2 and ZO-1 distribution. Scale bar: 100µm\ (E) Image of DAPI-stained (blue) section of the chick forebrain electroporated with the control construct (pCAG-IRES-GFP), where green fluorescence demarcates the domain of electroporation. Scale bar: 100µm (E′) Image of a DAPI-stained (blue) section of the chick forebrain electroporated with the control construct (pCAG-IRES-GFP), with ZO1 immunohistochemistry (red) (F) Image of DAPI-stained (blue) section of the chick forebrain electroporated with the Cdh-2 LOF construct (pCAG-DN-Cdh-2), where green fluorescence demarcates the domain of electroporation. Scale bar: 100µm (F′) Image of a DAPI-stained (blue) section of the chick forebrain electroporated with the Cdh-2 LOF construct (pCAG-DN-Cdh-2), with ZO1 immunohistochemistry (red). White stars indicate regions with loss of ZO1. (G) Image of DAPI-stained (blue) section of the chick forebrain electroporated with the Cdh-2 GOF construct (pCAG-Cdh-2-FL), where green fluorescence demarcates the domain of electroporation. Scale bar: 100µm (G′) Image of a DAPI-stained (blue) section of the chick forebrain electroporated with the Cdh-2 GOF construct (pCAG-Cdh-2-FL), with ZO1 immunohistochemistry (red). White stars indicate region with loss of ZO1. Scale bar = 100 μm

### Dynamic F-actin accumulation in the cortex of neuroepithelial cells mediated by Cdh-2 underlies spatiotemporal variability in roof plate stiffness

We observed that functional perturbation of Cdh-2 either through LOF or GOF resulted in ZO-1 depletion. However, ZO-1 is only present in apical adherens junctions; hence, the effects of its depletion should only be limited to the apical side. However, perturbation of Cdh-2 alters the stiffness of the entire dorsal forebrain; hence, this raises the question of how mechanical cues are transmitted across the entire length of the cell along the apicobasal axis. ZO-1 and Cdh-2 cooperate to establish and maintain apical adherens junctions by physically linking the cadherin–catenin complex to the tight junction scaffold and underlying actin cytoskeleton.

Myosin (non-muscle myosin II or NMII) contractility is well known to confer forces necessary to trigger mechanical change. Myosin II consists of heavy chains and regulatory light chains (MLC or MRLC). In its basal state, myosin exhibits low ATPase activity and limited ability to form bipolar filaments or generate force. Phosphorylation of the regulatory light chain activates myosin II. This phosphorylation enhances its interaction with actin, leading to stronger contractility of the actomyosin cortex. Consequently, this increases cortical tension and contributes to tissue-level mechanical properties such as stiffness (Handorf et al., 2015). In fact, actomyosin-generated tension fluctuations were shown to drive local tissue fluidization in the presomitic mesoderm in Cdh-2 mutant zebrafish embryos (Mongera et al., 2018).

To investigate the role of myosin contractility if any in the process of roof plate invagiantion, we first examined the expression of phosphorylated myosin (pMLC) at HH23. We found that pMLC was uniformly distributed along the mediolateral axis; although, it was restricted to apical junctions (Fig. S10A, A’). We then pharmacologically inhibited myosin light chain kinase (MLCK) using ML-7 to reduce actomyosin contractility (Cheng et al., 2015; Flatman, 2002; Lin et al., 2012). To validate the efficacy of ML-7 in vitro, we treated DF-1 cells with the inhibitor ML-7 and assessed pMLC expression via immunostaining (Fig. S10B, B’, C, C’, D, D’ E). Notably, ML-7 treatment did not disrupt the morphology of roof plate invagination (Fig. S10F, G) (Fig. S10F, G). Consistent with this, we found no significant changes in pMLC levels or distribution following Cdh2 manipulation (Fig. S10H, I, J). Since myosin contractility had no apparent influence on forebrain roof plate invagination, we next investigated whether there are any changes in the actin cytoskeleton itself in the roof plate.

Yu et al. proposed a mechanistic model wherein a positive feedback loop at cell-cell junctions involving cadherin causes trans-dimerization-induced local actin polymerization and actin tethering-induced cadherin immobilization and accumulation (Yu et al., 2022). This process directs the formation of a uniform actin mesh beneath the cell membrane (cell cortex). Inspired by this model, we examined the distribution of F-actin by performing phalloidin staining of neuroepithelial cells following Cdh-2 perturbation. We quantified the intensity of phalloidin staining up to 80 µm distance into the cell, both from the apical and basal sides of the neuroepithelium in sections of the forebrain roof plate (Fig. S11A, B and C). Since phalloidin binds to F-actin, the intensity of phalloidin staining can serve as a measure of the amount of F-actin. Upon perturbation of Cdh-2, the following aberrant changes in F-actin levels were evident at HH23: LOF of Cdh-2 led to a significant increase in accumulation of F-actin on the apical side, whereas GOF led to a symmetric increase in F-actin across the entire apicobasal axis in the midline apex and vortex regions, with no change in F-actin distribution and levels in the dorsolateral neuroepithelial cells (Fig. 6G, H, and I). The increased accumulation of apical F-actin at HH23 correlated with the increase in stiffness observed in the LOF of Cdh-2.

**Fig. 6.**
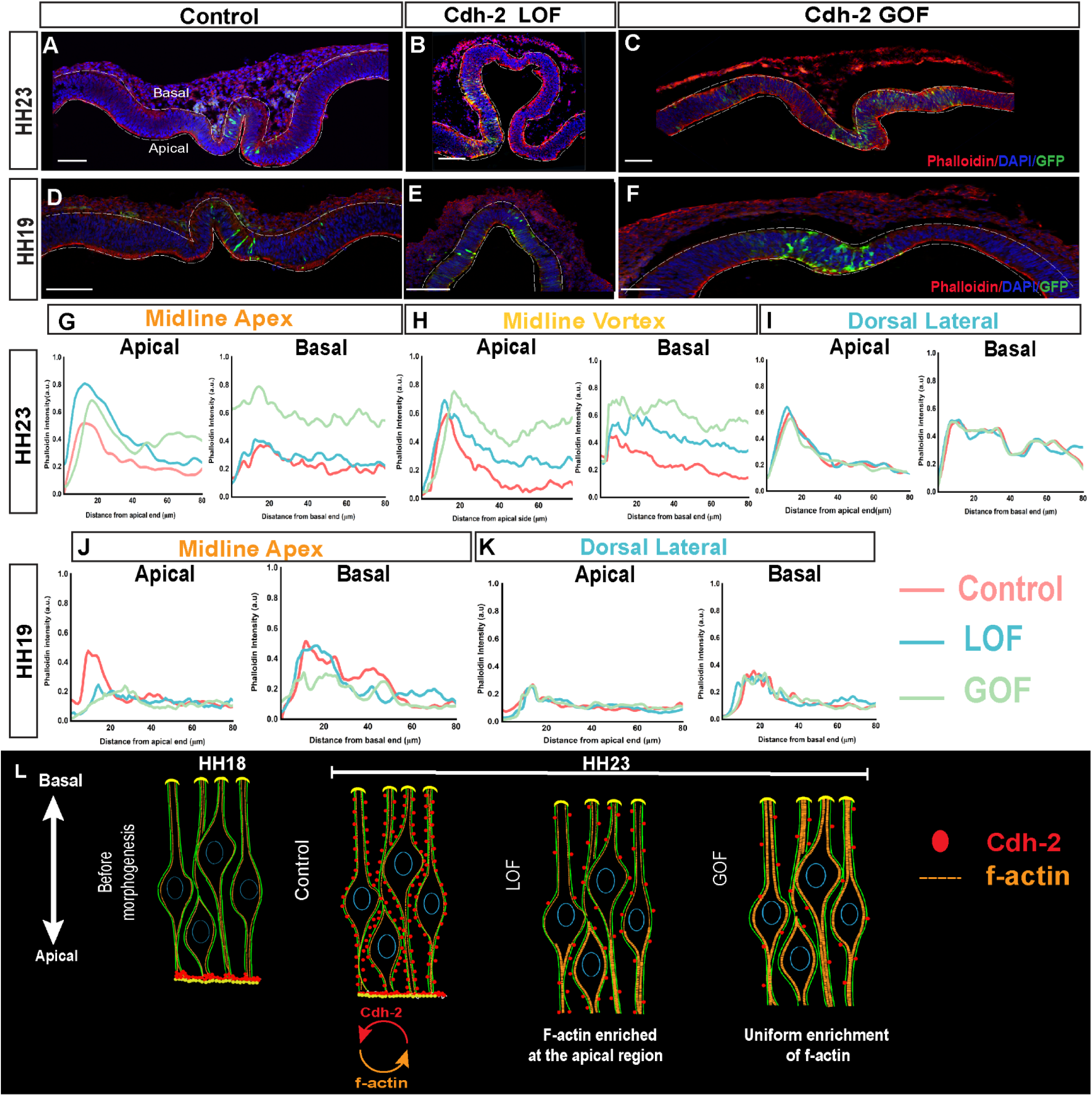
Cdh-2 regulates the spatial distribution of cortical F-actin in the roof plate neuroepithelial cells. (A) The merged image of a section of the control (pCAG-GFP) electroporated dorsal forebrain region at HH23, with DAPI (blue) marking the nuclei of neuroepithelial cells, phalloidin (red) marking the cortical actin within cells, and green marking the electroporated cells. Scale bar: 100 μm. (B) The merged image of a section of the Cdh-2 LOF (pCAG-DN-Cdh-2) electroporated dorsal forebrain region at HH23, with DAPI (blue) marking the nuclei of neuroepithelial cells, phalloidin (red) marking the cortical actin within cells, and green marking the electroporated cells. Scale bar: 100 μm. (C) The merged image of a section of the Cdh-2 GOF (pCAG-Cdh-2-FL) electroporated dorsal forebrain region at HH23, with DAPI (blue) marking the nuclei of neuroepithelial cells, phalloidin (red) marking the cortical actin within cells, and green marking the electroporated cells. Scale bar: 100 μm. (D) The merged image of a section of the control (pCAG-GFP) electroporated dorsal forebrain region at HH19, with DAPI (blue) marking the nuclei of neuroepithelial cells, phalloidin (red) marking the cortical actin within cells, and green marking the electroporated cells. Scale bar: 100 μm. (E) The merged image of a section of the Cdh-2 LOF (pCAG-DN-Cdh-2) electroporated dorsal forebrain region at HH19, with DAPI (blue) marking the nuclei of neuroepithelial cells, phalloidin (red) marking the cortical actin within cells, and green marking the electroporated cells. Scale bar: 100 μm. (F) The merged image of a section of the GOF of Cdh-2 (pCAG-Cdh-2-FL) electroporated dorsal forebrain region at HH19, with DAPI (blue) marking the nuclei of neuroepithelial cells, phalloidin (red) marking the cortical actin within cells, and green marking the electroporated cells. Scale bar: 100 μm (G,H,I) Phalloidin intensity profile of the neuroepithelium from the apical and basal surfaces from forebrains electroporated with constructs for control, LOF, and GOF of Cdh-2, at the midline apex (G), midline vortex (H), and dorsal lateral regions (I), at HH23. The fluorescence intensity of phalloidin was measured up to 80 microns into the cells from both the apical and basal surfaces. Intensity was normalised to a range of 0 to 1 by dividing it by the highest intensity, which was 85 for the control and all test groups. Control is shown in red, Cdh-2 LOF in cyan, and Cdh-2 GOF in green in all the plots. For every N, 30 lines of 80 AU were drawn, and mean value was calculated for each N. (J, K) Phalloidin intensity profile of the neuroepithelium from the apical and basal surfaces from forebrains electroporated with constructs for control, LOF, and GOF of Cdh-2, at the midline (J) and dorsal lateral regions (K), at HH19. The fluorescence intensity of phalloidin was measured up to 80 microns into the cells from both the apical and basal surfaces. Intensity was normalised to a range of 0 to 1 by dividing it by the highest intensity, which was 85 for the control and all test groups. Control is shown in red, Cdh-2 LOF in cyan, and Cdh-2 GOF in green in all the plots. For every N, 30 lines of 80 AU were drawn, and mean value was calculated for each N. (L) A schematic of the neuroepithelial cells from the dorsal forebrain region, showing Cdh-2 distribution (red circles) and cortical F-actin levels (yellow dashed lines) in the unmanipulated control forebrain at HH18 (prior to onset of morphogenesis) and at HH23 (W-shaped invagination formed). Further changes in Cdh-2 distribution (red circles) and cortical F-actin levels (yellow dashed lines) are shown when Cdh-2 is perturbed through LOF and GOF manipulations.

We also measured F-actin accumulation in the neuroepithelial cells at HH19, the stage at which the “pucker” formation was observed in the roof plate midline. In the control, in the midline apex region where the “pucker” is present, F-actin accumulation was significantly high in the apical as well as the basal sides of the neuroepithelium, while it was uniformly low across the apico-basal axis in the dorsolateral regions. However, both LOF and GOF of Cdh-2 led to significant changes in F-actin accumulation along the apicobasal axis compared with the control, particularly in the midline region (Fig. 6J and K). Specifically, the LOF of Cdh-2 reduced F-actin accumulation significantly on the apical side of the midline neuroepithelium, while GOF led to a symmetric decrease in F-actin accumulation across the apicobasal axis in the midline regions (Fig. 6L). These observations demonstrate that Cdh-2 perturbation disrupts the cortical F-actin distribution along the apicobasal axis of neuroepithelial cells.

The concentration of Cdh-2 at the cell membrane of neuroepithelial cells determines the adhesivity between them and generates adhesive force (adhesion tension) throughout the tissue. In contrast, cortical F-actin accumulation influences cellular morphology, such that increased F-actin levels reduce cellular flexibility and generate a counteracting force, that is, cortical tension. The balance between Cdh-2-mediated adhesion and F-actin-generated cortical tension determines precise cellular organization within a tissue. In this regulatory framework, the modulation of Cdh-2 expression directly influences the pattern of F-actin accumulation. Furthermore, Cdh-2 expression directly modulates the stiffness of the dorsal forebrain. We hypothesized that the precise spatiotemporal accumulation of Cdh-2 at the cell surface and F-actin in the cortex determines neuroepithelial stiffness. The establishment of a normal stiffness gradient is critical for the formation of a pucker at HH19. Upon Cdh-2 loss-of-function (LOF), increased tissue stiffness resulting from biased cortical F-actin accumulation prevents the roof plate from achieving sufficient flexibility to form the characteristic W-shaped morphology. Instead, uniformly high levels of stiffness across the dorsal forebrain lead to evagination. In contrast, the gain-of-function (GOF) of Cdh-2 leads to uniform actin accumulation, disrupting the normal apicobasal F-actin distribution required for localized flexibility and proper invagination.

## Discussion

Morphogenesis is orchestrated by the interplay between mechanical forces and molecular signaling. Until recently, most reports have focused on forces generated by pMLC contractility, which promote actin polymerization and drive cortical rearrangements and cell shape changes. The present study establishes a mechanistic framework that links cell adhesion dynamics, actin cytoskeleton remodeling to modulate cell-cell rearrangement dynamics required for flexibility of the tissue during morphogenesis.

In the chick embryo, at HH18, the dorsal forebrain neuroepithelium exhibits a relatively uniform stiffness of approximately 800 kPa. By HH21, this tissue undergoes morphological changes accompanied by the emergence of a graded stiffness landscape: the midline apex becomes significantly more compliant (∼67 kPa), while the dorsolateral regions maintain a higher stiffness of approximately 400 kPa. The progressive formation of this ∼10-fold stiffness gradient preceeds the transformation of the relatively flat roof plate neuroepithelium into a complex W-shaped invagination. This mechanical patterning is consistent with established principles of epithelial morphogenesis; whereby differential mechanical properties generate the physical conditions necessary for controlled tissue deformations.

We observed that the overlying mesenchyme initially maintains consistently high stiffness, with a marked increase to ∼750 kPa specifically in the region directly above the midline by HH23. This change just precedes the formation of the W-shaped invagination. The increasing stiffness of the overlying mesenchyme imposes a mechanical constraint that directs the more compliant neuroepithelium to buckle inward rather than expand outward. Such interlayer mechanical coupling is recognized as a fundamental mechanism in neural tube closure and other morphogenetic processes (Christodoulou & Skourides, 2023). We found that progressive enrichment of Cdh-2 protein along the apicobasal axis of the neuroepithelial cells correlates with the evolving tissue stiffness and morphological changes in the dorsal forebrain. Loss-of-function of Cdh-2 results in uniformly elevated stiffness across the roof plate and leads to tissue evagination, whereas gain-of-function produces a V-shaped invagination with altered mechanical properties. Notably, both perturbations abolish the normal stiffness gradient, demonstrating that precise spatial and temporal regulation of Cdh-2 is essential for changes in stiffness which are likely to set the stage for proper morphogenesis.

Quantitative analysis of Cdh-2 levels at HH23 following overexpression suggests activation of a negative feedback mechanism that maintains cell-surface Cdh-2 during invagination of the forebrain roof plate. This mechanism ensures optimal distribution of Cdh-2 along the apicobasal axis in the neuroepithelium, essential for robust morphogenesis. Early perturbation of Cdh-2 disrupts this progressive increase, resulting in undetectable Cdh-2 at HH19 and significantly reduced levels at HH23. We have previously reported that dorsal forebrain exhibits differential proliferation in the neuroepithelium (Gupta & Sen, 2015; Udaykumar et al., 2023). It is likely that proliferating cells in the dorsolateral neuroepithelium generate forces directed toward the midline of the roof plate which has very little cell proliferation. The present study confirms that these proliferation derived forces appear to be independent of Cdh-2 activity and may act in parallel to support the observed morphological changes. Adherens junction scaffolding protein ZO-1 is one of the mediators linking Cdh-2 to the cytoskeletal machinery. We found that both loss- and gain-of-function of Cdh-2 resulted in depletion of ZO-1, indicating localized loss of junctional integrity. This indicates that ZO-1 couples cadherin–catenin complexes to the actin cytoskeleton which may function by modulating mechanical tension at apical adherens junctions. A similar role of ZO-1 was highlighted in another study where it was found to regulate tension on VE-cadherin-based junctions and orchestrates spatial actomyosin organization (Tornavaca et al., 2015).

The apicobasal extension of Cdh-2 expression during roof plate morphogenesis indicates active regulatory dynamics beyond apical junctions. We therefore examined actomyosin dynamics across the dorsal forebrain neuroepithelium and found that phosphorylated myosin remained largely restricted to junctional regions and was unaffected by Cdh-2 perturbations. In contrast, Cdh-2 manipulations markedly altered the distribution and accumulation of cortical F-actin along the apicobasal axis in neuroepithelial cells.

These findings align with established models in which cadherin-mediated adhesion regulates cortical actin organization. As elegantly demonstrated by Yu et al., surface recruitment of cadherins not only promotes their own clustering but also drives the recruitment and stabilization of cortical F-actin through a positive feedback mechanism. Loss of ZO1 from apical junctions initiates the disassembly of apical adherens junctions, which in turn disrupts the F-actin–Cdh2 positive feedback loop across the cell cortex. This disruption propagates like a domino effect, spreading not only along the apico-basal axis but also to neighboring cells. In the roof plate neuroepithelium, active recruitment of Cdh-2 to the cell surface thus generates dynamic adhesion forces that, in turn, modulate cortical actin dynamics. The resulting “tug-of-war” between Cdh-2 mediated adhesion tension at the cell surface and cortical tension in the dynamic cell cortex generates a residual force that dictates the mechanical flexibility of the neuroepithelium. This enables coordinated bending in response to tangential expansion from highly proliferative neighboring cells and compressive forces from the overlying mesenchyme.

Paradoxically, loss of Cdh-2 led to enhanced apical F-actin accumulation at HH23. This is likely to arise from compensatory cortical remodeling triggered by insufficient Cdh-2 levels. In the absence of optimal Cdh-2, the cell cortex undergoes adaptive reorganization to retain cell-cell adhesion to maintain tissue integrity while perturbing tissue stiffness required. This interpretation is supported by previous reports showing that cortical actin organization rapidly readjusts to preserve tissue integrity upon perturbation (Sakamoto et al., 2023).

Collectively, our findings reveal an active mechanochemical feedback loop mediated by Cdh-2 that adds to the forces essential for neuroepithelial bending into a W-shape against the stiff overlying mesenchyme. The study highlights that the forces driving morphogenesis are not derived solely from myosin contractility; rather, they can be generated by adhesion dynamics. In the developing forebrain, Cdh-2 functions not merely as a passive cell adhesion molecule but as an active regulator that imparts specific mechanical properties to the tissue.

This work advances our understanding of how cells within a tissue collectively generate and respond to mechanical forces by utilizing molecular players on the cell surface, rather than relying exclusively on energy-consuming motors to determine the tissue’s mechanical properties. Furthermore, this study underscores the importance of accurately defining tissue stiffness in biological systems. We have demonstrated that, in addition to cortical tension, adhesion tension plays a significant role. Other forces are also at play in the roof plate, including tangential forces and possibly pressure from cerebrospinal fluid (CSF). Nevertheless, we excluded tangential forces as a direct driver of tissue stiffness gradients. Consistent with this, previous literature has established that eCSF pressure primarily drives global tangential growth and vesicle expansion, rather than directly mediating chick roof plate morphogenesis. (K. E. Garcia et al., 2019). Another key force is likely to be the compressive force exerted by the overlying mesenchyme. Therefore, the effective stiffness of the roof plate represents the combined influence of all these forces during its morphogenesis. Future studies combining live force measurements with molecular dissection of Cdh-2-associated signaling complexes will be essential to fully elucidate the regulation of tissue mechanics during morphogenesis. In addition, investigating other junctional components and mesenchymal factors will help clarify the multi-tissue mechanical interactions underlying epithelial folding.

Finally, these findings have important implications for understanding holoprosencephaly (HPE). While genetic mutations in signaling pathways (e.g., Sonic hedgehog) are well-established contributors to HPE, the role of mechanical forces has received comparatively less attention. Our demonstration that Cdh-2-mediated mechanical regulation is essential for proper forebrain roof plate invagination suggests that defects in mechanotransduction pathways may also contribute to HPE pathogenesis.

## Materials and Methods

### Chicken Eggs and Embryo Handling

Day old chicks of CARI PRIYA strain of the White Leghorn variety of chickens were procured from the Central Avian Research Institute, Izatnagar, Bareilly, Uttar Pradesh, India. These birds were maintained by Ganesh Enterprises in Nankari Village, Kanpur, Uttar Pradesh, India, and fertilized chicken eggs were procured from them. Eggs were incubated at 38 °C in a humidified incubator until the embryos reached the desired developmental stages, which was determined using the Hamburger and Hamilton staging system. Approval for conducting experiments with the fertilized chicken eggs was granted by the Animal Ethics Committee, IIT Kanpur. Animal husbandry, supply, maintenance, and care in the animal facility before and during the experiments fully met the needs and welfare of animals.

### Cell culture

For the sensor assay, human embryonic kidney fibroblast cells (HEK293T) (ATCC, CRL3216) were transfected using Turbofect, according to the manufacturer’s protocol.

### Tissue preparation for immunohistochemistry and RNA in situ hybridization

Heads were collected from both unmanipulated and electroporated embryos at the specified stages and fixed overnight in 4% paraformaldehyde (Catalog no. P6148, Sigma Aldrich). For cryoprotection, the samples were subjected to sucrose gradient immersion, first in 15% and then 30% sucrose solutions prepared in phosphate-buffered saline (PBS). Post-cryoprotection, the tissues were embedded in Polyfreeze/OCT compounds) and sectioned coronally at a thickness of 10 μm using a Leica CM1520 cryostat.

### Tissue Preparation for AFM

Chick brain tissues were collected at appropriate developmental stages in phosphate-buffered saline (PBS). The tissues were then treated with 15% sucrose in 1X PBS for 4 h at 4°C, followed by treatment with 30% sucrose for 4 h at room temperature (RT). Afterward, the brains were embedded in Optimal Cutting Temperature (OCT/Polyfreeze) compound (Catalog no. P0019, Sigma Aldrich) and stored at −80°C. Cryosectioning was performed using a cryostat (Leica CM1850). Coronal sections of 2 μm thickness were placed on coverslips (Blue-Star, 22 × 22 mm) pre-coated with 0.1% poly-L-lysine solution.

The chick forebrain is double-layered and, at these embryonic stages, is viscid and friable. Consequently, probes in contact mode tend to become entrapped on the surface. We therefore used sections as thin as 2 μm and maintained an indentation depth not exceeding 4 μm. Because we worked with tissue sections, we attempted to span the entire roof plate in the mid-posterior region and performed atomic force microscopy (AFM) on multiple sections, averaging the readings.

### Plasmids

**pCAG-dn-Cdh-2-IRES-GFP:** The sequence of DNA provided below including EcoRI and NotI restriction sites was synthesized (Macrogen), followed by digestion with EcoRI and NotI restriction enzymes. To generate the CAG-IRES-GFP vector, the pCAG-NeuroD1-IRES-GFP plasmid (Add gene, 45025) was digested with EcoRI and NotI. The resulting restriction-digested synthesized DNA frag was ligated with the digested vector using T4 DNA ligase, leading to the formation of pCAG-dn-Cdh-2-IRES-GFP. This was inspired by the dominant negative Cdh-2 construct described in literature (Kintner, 1992).

### Cdh-2 without extracellular domain (DNA)

5’ATGTGCCGGATAGCGGGAACGCCGCCGCGGATCCTGCCGCCGCTGGCGCTGATGCTGCTGGCGGCCCTGCAGCAGGCACCGATAAAAGCAACTTGTGAAGACATGTTGTGCAAGATGGGATTTCCTGAAGATGTGCACAGTGCAGTCGTGTCGAGGAGTGTACATGGAGGACAACCTCTGCTCAATGTGAGGTTTCAAAGCTGCGATGAAAACAGAAAAATATACTTTGGAAGCAGTGAGCCAGAAGATTTTAGAGTAGGTGAAGATGGTGTGGTATATGCAGAGAGAAGCTTTCAACTTTCAGCAGAGCCCACGGAGTTTGTAGTGTCTGCTCGAGACAAGGAAACTCAGGAAGAATGGCAAATGAAGGTGAAGCTACTGCAGCCTAATGCTATTAACATCACTGCTGTAGACCCTGACATTGATCCAAATGCAGGCCCATTTGCCTTTGAGCTGCCTGATTCACCTCCTAGTATTAAGAGGAATTGGACCATTGTTCGAATTAGTGGTGATCATGCCCAGCTCTCTTTAAGGATCAGGTTCCTGGAGGCTGGTATCTATGATGTGCCCATAGTAATTACAGATTCTGGAAATCCACATGCATCTAGCACTTCTGTGCTAAAAGTGAAAGTTTGCCAATGTGACATAAATGGGAGACTGTACTGATGTTGACCGGATTGTTGGCGCAGGACTGGGCACTGGTGCCATCATTGCAATTCTGCTTTGTATCATCATCTTACTCATTTTAGTTTTGATGTTCGTAGTATGGATGAAGCGCCGTGATAAGGAGCGTCAGGCCAAGCAGCTCTTAATTGATCCAGAAGATGATGTGAGGGACAACATTCTGAAATATGATGAAGAAGGTGGTGGAGAAGAAGATCAGGATTATGACTTGAGCCAGCTCCAGCAGCCTGCACTGTAGAACCAGACGCCATCAAACCTGTTGGAATCAGACGTCTTGATGAAAGGCCAATCCATGCAGAACCTCAGTATCCAGTCAGATCAGCTGCTCCTCATCCTGGGGACATTGGGGACTTCATTAATGAGGGACTTAAAGCAGCCGACAACGACCCTACAGCCCCGCCATACGATTCCCTCTTAGTCTTTGACTATGAAGGAAGCGGCTCCACTGCTGGATCCTTGAGCTCTCTTAATTCCTCAAGTAGCGGTGGTGAGCAAGACTATGACTACCTAAATGACTGGGGCCCACGTTTCAAGAAACTTGCTGACATGTAGGTGGAGGTGATGACTGA3’

### Full length Cdh-2

This construction was provided by Prof. C. Cepko from Harvard Medical School, USA.

### Sensor Assay

To conduct the sensor assay, a Cdh-2-sensor construct was created and designated as pCAG a mCherry-Cdh-2-sensor. The 3’ UTR of the chick Cdh-2 gene, which includes the target sequence for two Cdh-2 miRNAs, was PCR-amplified using the following primers:

Forward primer:

5′ATAGCGGCCGCTAGCACTTCAAAGTGAACTTTGTTTCTGG3′ (Not I)

Reverse primer:

5′GGCAAGCTTGTAGTCGACAATTTTCAGTCTCCTTATTTTAATAAAAGC3′(HindIII)

PCR amplification was performed using Phusion polymerase. The resulting amplified product was digested with NotI and HindIII restriction enzymes. This digested product was ligated into the pCAG-mCherry vector, which was also digested similarly. For the sensor assay, pCAG-mCherry-LacZ served as a negative control, as this construct did not contain Cdh-2 RNAi target sequences. Transfection included either pCAG-mCherry-Cdh-2-sensor or pCAG-mCherry-LacZ, alone or in combination with pRmiR-Cdh-2-RNAi-Oligo1 or pRmiR-Cdh-2-RNAi-Oligo2 at a molar ratio of 6:1. The transfection was done with HEK293T cells at 70% confluency. The expression of GFP or mCherry in transfected cells was subsequently observed. The mean fluorescence intensity was quantified using ImageJ software. An unpaired t-test was conducted using Origin Pro 2024b software to calculate the mean of all replicates, along with standard deviation (SD), standard error of the mean (SEM), and p-values.

### In-ovo electroporation

After 24 h of incubation, 3 ml of albumin was carefully removed to lower the embryo position inside the egg. A small window was then created on the eggshell above the forebrain vesicle at Hamburger-Hamilton stage 10 (HH10). DNA constructs (300ng to 1 µg/ml) mixed with 0.1% Fast Green (Catalog no. F7258, Sigma Aldrich) for visualization were injected into the forebrain vesicle. Electroporation was performed using platinum “hockey stick” electrodes (NepaGene, CUY611P3-1), positioned such that the positive electrode rested over the dorsal forebrain and the negative electrode beneath the yolk. Five 15 V square pulses, each lasting 50 ms, were applied at 950 ms intervals, utilizing a BTX ECM830 electroporator (450662). After electroporation, sterile PBS containing penicillin and streptomycin was gently added to the embryos and the egg windows were sealed with tape. The eggs were returned to the incubator until the embryos reached the desired stages for tissue collection.

### Injections

After 24 h of incubation, 3 ml of albumin was carefully removed to lower the embryo position inside the egg. A small window was then created on the eggshell above the forebrain vesicle at Hamburger-Hamilton stage 10 (HH10). ML-7, Myosin light chain kinase inhibitor (Catalog no. I2764, Sigma-Aldrich) in DMSO (50 µM) mixed with 0.1% Fast Green (Catalog no. F7258, Sigma Aldrich) for visualization were injected into the forebrain vesicle. After injections, sterile PBS containing penicillin and streptomycin was gently added to the embryos and the egg windows were sealed with tape. The eggs were returned to the incubator until the embryos reached the desired stages for tissue collection.

### RNA in situ hybridization

Complementary DNA (cDNA) clones were used to generate digoxigenin labelled riboprobes for mRNA in situ hybridization. Antisense riboprobes were synthesized using appropriate RNA polymerases, such as T3 RNA polymerase (Catalog no. P2083, Promega), T7 RNA polymerase (Catalog no. P2075, Promega) or SP6 RNA polymerase (Catalog no. P4084, Promega). RNA in situ hybridization was conducted on coronal forebrain sections, following established methods (Trimarchi et al., 2007). Digoxigenin-labelled riboprobes for hybridization were synthesized via in vitro transcription of previously described cDNA clones.

### Immunohistochemistry

Immunostaining of cryosections mounted on Poly-L-Lysine (Catalog no. P8920, Sigma Aldrich) coated slides involved three washes in PBS with 0.1% Tween-20 (PBT) for 10 min in PFA, followed by a 45 min block with 10% heat-inactivated goat serum in PBT. Primary antibodies were added to the blocking solution and incubated overnight at 4 °C. After washing three times with PBT for 5 min each, secondary antibodies (1:250) were incubated for 2 h at room temperature. Antibody used were Anti-N-Cadherin (Catalog no. C3865, Sigma-Aldrich), Anti-GFP (Catalog no. A-6455, ThermoFisher), Anti-phospho-Histone 3 (Catalog no. H0412, Sigma-Aldrich), Anti Zona occludens-1 (Catalog no. 61-7300, Invitrogen), Phospho-Myosin Light Chain 2 -Thr18/Ser19 (Catalog no. 3674, CST), Alexa Fluor 488-conjugated anti-mouse IgG (Catalog no.115-545-003, Jackson ImmunoResearch Labs), Alexa Fluor 594-conjugated anti-mouse IgG (Catalog no. 115-585-003, Jackson ImmunoResearch Labs), Alexa Fluor 594-conjugated anti-rabbit IgG (Catalog no.111-585-045, Jackson ImmunoResearch Labs), Alexa Fluor 488-conjugated anti-rabbit IgG (Catalog no. 111-545-003, Jackson ImmunoResearch Labs), Phalloidin conjugated to Alexa Fluor 647 (Catalog no. A22287, ThermoFisher). For visualization of F-actin, the sections were incubated with phalloidin for 45 min at room temperature with 1% BSA (Hi Media MB083) in PBS. All sections were subsequently stained with DAPI (Catalog no. D9542-10MG, Sigma Aldrich) to mark nucleus.

### Immunohistochemistry on DF1 cells

DF-1 cells were seeded onto gelatin-coated coverslips to achieve 80% confluency at the time of transfection. Twenty-four hours post-seeding, the cells were treated with either DMSO or ML-7. Following a 3- or 12-hour incubation, the cells were immunoassayed according to the previously described protocol. The coverslips were washed three times with PBT for 5 minutes each, counterstained with DAPI, and mounted using an antifade medium.

### Image Acquisition

Fluorescent images were captured using an AXR Nikon confocal microscope and a Carl Zeiss LSM780NLO multiphoton laser scanning confocal microscope, with accompanying software NIS Elements AR 5.42.04 and ZEN 2012 for image processing. RNA in situ hybridization images were obtained using a Leica DM500B stereomicroscope equipped with a DFC500 camera. The final image assembly and figure preparation were performed using Adobe Photoshop and Adobe Illustrator 2024.

### Replicate definition

The experiments were performed with biological and technical replicates, as indicated. A biological replicate corresponded to one chicken embryo used for the experiment. Experiments with at least three biological replicates were used to calculate the statistical significance for each analysis. All graphs represent the mean ± SEM. All statistical analyses were performed, and graphs were generated using OriginPro software.

### Quantification of Youngs Moduli/Elastic Moduli for stiffness measurements

To calculate Young’s modulus, an atomic force microscope (Oxford Instruments MFP3D origin) was used. Heat maps were generated with the help of Ms. Deepali Ubale, a technician at the Nanoscience Department of IIT Kanpur, India. Asylum Research Software was used to calculate Young’s modulus. The Hertz model was used to generate heat maps. The probe used was from Oxford Instruments silicon probe with Au coating and a frequency of 100KHz was used. Spring Constant ≥ 0.3. The applied force was 10nN and the probe size was 10 nm. The Hertz model outlines the deformation characteristics of purely linearly elastic materials (Johnson, 1995). Therefore, when fitting any linear elastic data segment to the Hertz model for the modulus of elasticity (E), the resulting E will remain consistent across other linear elastic segments. Based on the comprehensive literature and derivation, the final Hertz equation for a spherical AFM tip is

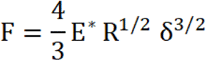

where:

F = Applied force (N)

E* = Reduced elastic modulus (Pa)

R = Radius of the spherical AFM tip (m)

δ = Indentation depth (m)

The reduced elastic modulus E* is defined as:

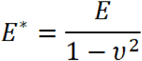

where:

E = Young’s modulus of the sample (Pa)

ν = Poisson’s ratio of the sample

After generating the heat maps, we selected those with uniform colors and averaged them to determine the Young’s modulus for 100 indentation points. For this analysis, three blocks (20 × 20 μm²) were scanned, each containing 100 points, with a focus on a specific region. This process was repeated for 6–8 embryos. Since tissue sections were embedded in OCT compound, we measured the stiffness of OCT alone without the embedded tissue, which yielded a value of 0.2 kPa. The relationship between OCT and tissue measurements is nonlinear. Consequently, we were unable to normalize the values to correct for the effects of tissue embedding in OCT. We therefore plotted the raw data obtained from the spatial heat maps. We therefore emphasized the trends in the data rather than the precise absolute values.

### Quantification of Phalloidin fluorescence intensity

Phalloidin fluorescence intensity was quantified from the images acquired at 60X magnification. For each section, three distinct regions of interest (ROIs) were marked: midline apex, midline lateral, and dorsal lateral regions. Within each ROI, 30 lines were drawn, extending 80 μm from both the apical and basal surfaces of neuroepithelial cells (indicated by red and green lines). Utilizing the ‘Plot Profile’ function in ImageJ software, fluorescence intensity profiles were generated along each line. The mean intensity values for each point across the lines are plotted for each ROI. To facilitate comparison, the intensity values were normalized to a range of 0–1 by dividing by 85 and plotted on the y-axis with the distance represented on the X-axis. This normalization was based on the maximum intensity value obtainable owing to the saturation limit of ImageJ, where 0 represents the lowest intensity and 1 represents the highest intensity. (N=3)

### Quantification of pH3 positive cells

A rectangle measuring 375 μm × 375 μm was overlaid on the image at 40X magnification. The numbers of pH3 positive (red), GFP-positive (green), and double-positive (yellow) cells were counted manually using the ImageJ software. N=5 for midline and N=4 for lateral measurements.

## Acknowledgments

The authors acknowledge Prof. Constance Cepko (Harvard Medical School, Boston, MA, USA) for the cDNA for Cdh-2 FL (chick). The authors acknowledge Deepali Ubale for help with atomic force microscopy and force mapping and the Department of Nanoscience at IIT Kanpur for the AFM facility. The authors acknowledge Ms. Neetu Dey for help with the multiphoton laser scanning confocal microscope. The authors acknowledge Ms. Rashmi Parihar for help with generation of 2 µm cryosection at the cryo-sectioning facility at CEAF, IIT Kanpur. The authors are grateful to Dr. Sandeep Gupta, Dr. Tathagata Biswas, and Prof. Amitabha Bandyopadhyay for critical comments on the manuscript. The authors acknowledge Mr. Naresh Gupta for technical support in this study.

## Author contributions

Conceptualization: M.S, J.S Methodology: M.S., P.S., P.G., S.K. M.A.A.Z. and J.S. Software: M.S. Formal Analysis: M.S, J.S. Investigation: M.S, J.S. Resources: J.S. Data Curation: M.S., J.S. Visualization: M.S., S.K., J.S. Supervision: J.S. Writing original draft: M.S., J.S. Writing & editing: M.S., P.G., J.S. Project administration: J.S. Funding acquisition: J.S.

## Data and materials availability

The construct expressing dominant negative CDH-2 and CDH-2 RNAi generated during this study will be shared upon coverage of shipping costs. The detailed protocol for measuring the stiffness of the embryonic forebrain tissues of *Gallus gallus* would also be shared upon request.

No standardized data types were generated in this study. All data supporting the conclusions can be found in the main text or are available upon request from the corresponding author. Any additional information required to reanalyze the data reported in this study is available from the lead contact upon request. Requests for further information and resources should be directed to and will be fulfilled by the lead contact, Jonaki Sen

