## Supplementary material for "Cdh-2 and cortical f-actin dynamically cooperate to establish a stiffness gradient which contributes to forebrain roof plate invagination": SUPPLEMETARY FIGURES AND LEGENDS

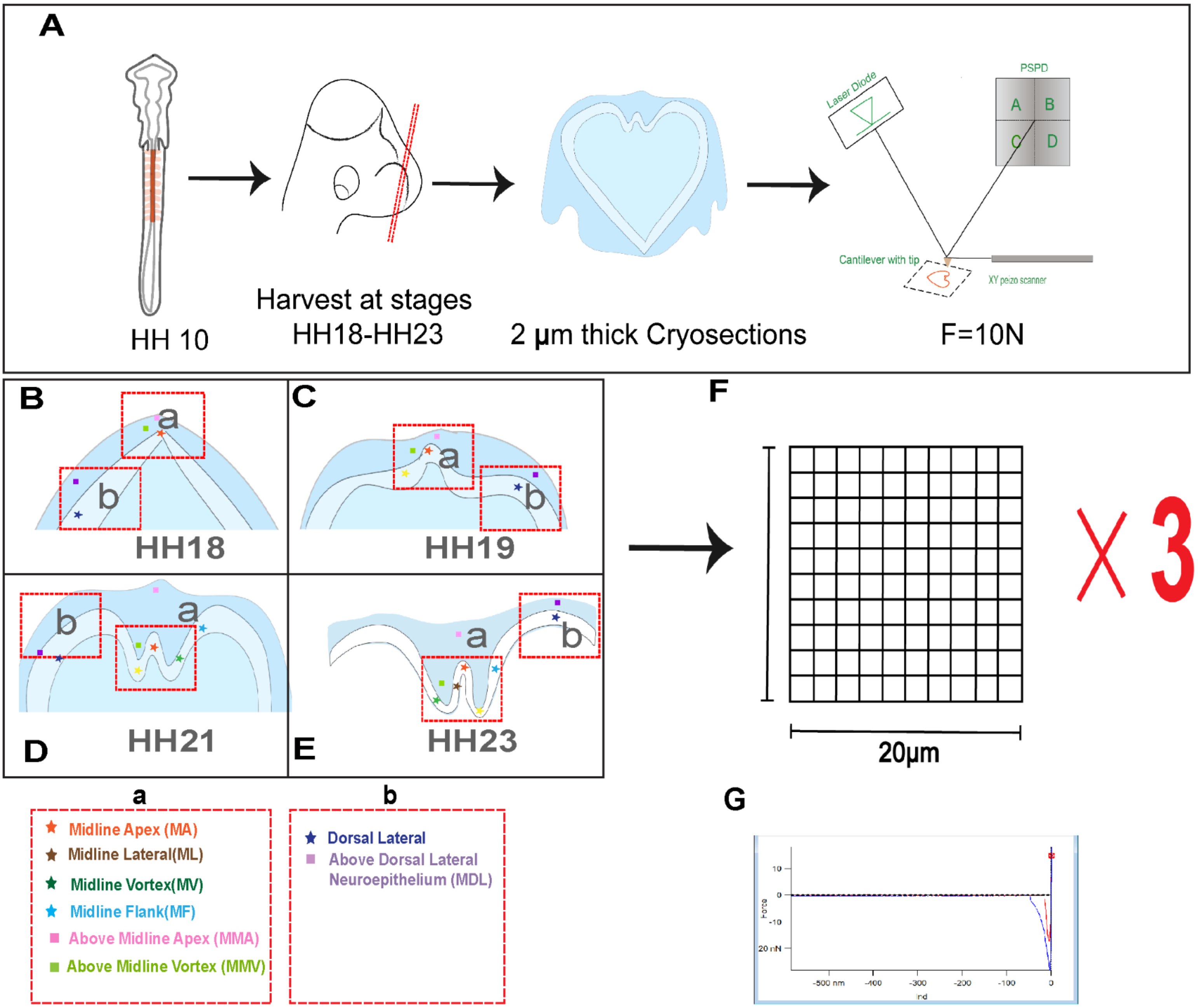


**Fig. S1. An optimized protocol for force mapping in the chick forebrain.**

(A) A schematic for the optimised protocol developed for measuring stiffness of the chick forebrain using atomic force microscopy (AFM) and an illustration of contact mode AFM whereby a finely pointed tip mounted on a cantilever is used to perform detailed scans of samples and a laser beam is used to sense the bending of the cantilever as its tip hits the surface. The Young's modulus values obtained are used to determine the stiffness of the material. For the dorsal forebrain tissues a precise force of 10 nN was applied to the cantilever and its resulting deflection was measured to determine the stiffness.

(B,C,D,E) Schematic of coronal sections of the dorsal forebrain at stages HH18, HH19, HH21 and HH23 with red-dashed outline boxes denoting the regions of the midline (a) and dorsolateral (b) tissues where AFM was used to measure stiffness. The various subdomains named MA, ML, MV, MF and DL within the neuroepithelium are marked with coloured stars. The various subdomains of the mesenchyme named MMV, MMA and MDL are marked by coloured squares.

(F) Schematic depicting the 20 X 20 $\mu m^2$ area within each subdomain which was further divided into 100 squares, and the 100 squares in three such 20 X 20 $\mu m^2$ were scanned to measure Young's modulus for each N.

(G) A representative Force vs Indentation Curve obtained after scanning with the AFM probe in each of the 100 squares shown in (F). 100 such graphs are combined to produce the heat map of one 20 X 20 $\mu m^2$ shown in Fig. S2.

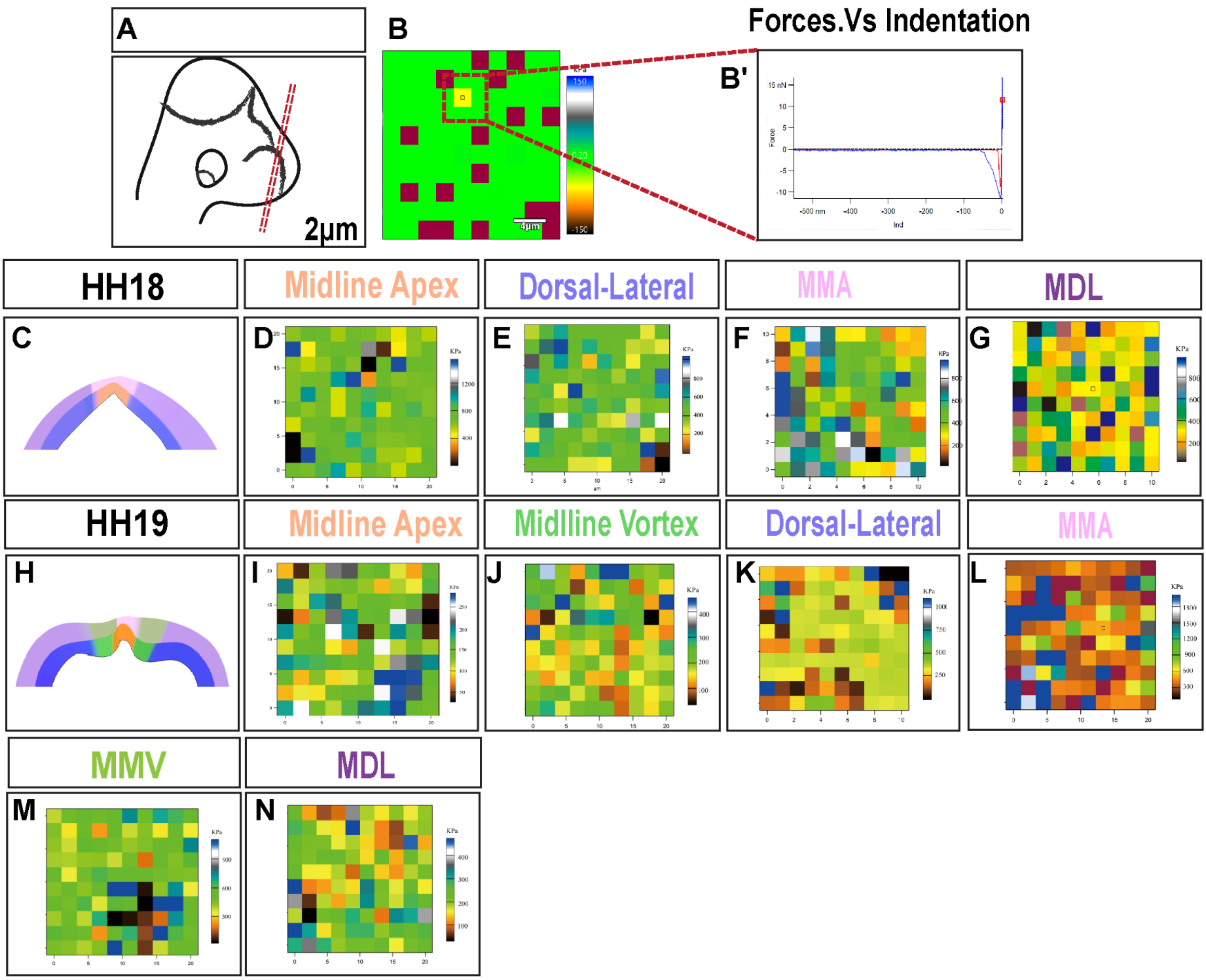


**Fig. S2. Heat maps for the measured Young's modulus in the chick forebrain at HH18 and HH19.**

(A) A schematic representation of the chick forebrain with the red dashed line demarcating plane in the middle posterior region from which 2 µm thick coronal slices were taken.

(B) Representative heat map of Young's Modulus (YM) measured for each of 100 squares within a 20*20 µm² area of each subdomain of the dorsal forebrain tissues (neuroepithelium and mesenchyme) that were scanned to measure YM in triplicate.

(B') Representative Force vs Indentation Curve for one representative square (in the red dashed box) out of the 100 square heat maps of 20*20 µm² area shown B.

(C) A schematic representation of a coronal slice from the middle posterior region of the chick dorsal forebrain at HH18 with distinct colours marking subdomains of the roof plate neuroepithelium and overlying mesenchyme.

(D, E and F, G) The representative heat maps for the subdomains of the neuroepithelium (midline apex and dorsal lateral) and the mesenchyme (MMA and MDL) at HH18. For each region, a 20*20 $\mu m^2$ area was scanned to measure YM in triplicate.

(H) A schematic representation of a coronal slice from the middle posterior region of the chick dorsal forebrain at HH19 with distinct colours marking subdomains of the roof plate neuroepithelium and overlying mesenchyme.

(I,J,K and L,M,N) The representative heat maps for the subdomains of the neuroepithelium (Midline Apex, Midline Vortex and Dorsal-Lateral) and the mesenchyme (MMA, MMV and MDL) at HH19. For each region, a 20*20 $\mu m^2$ area was scanned to measure YM in triplicate.

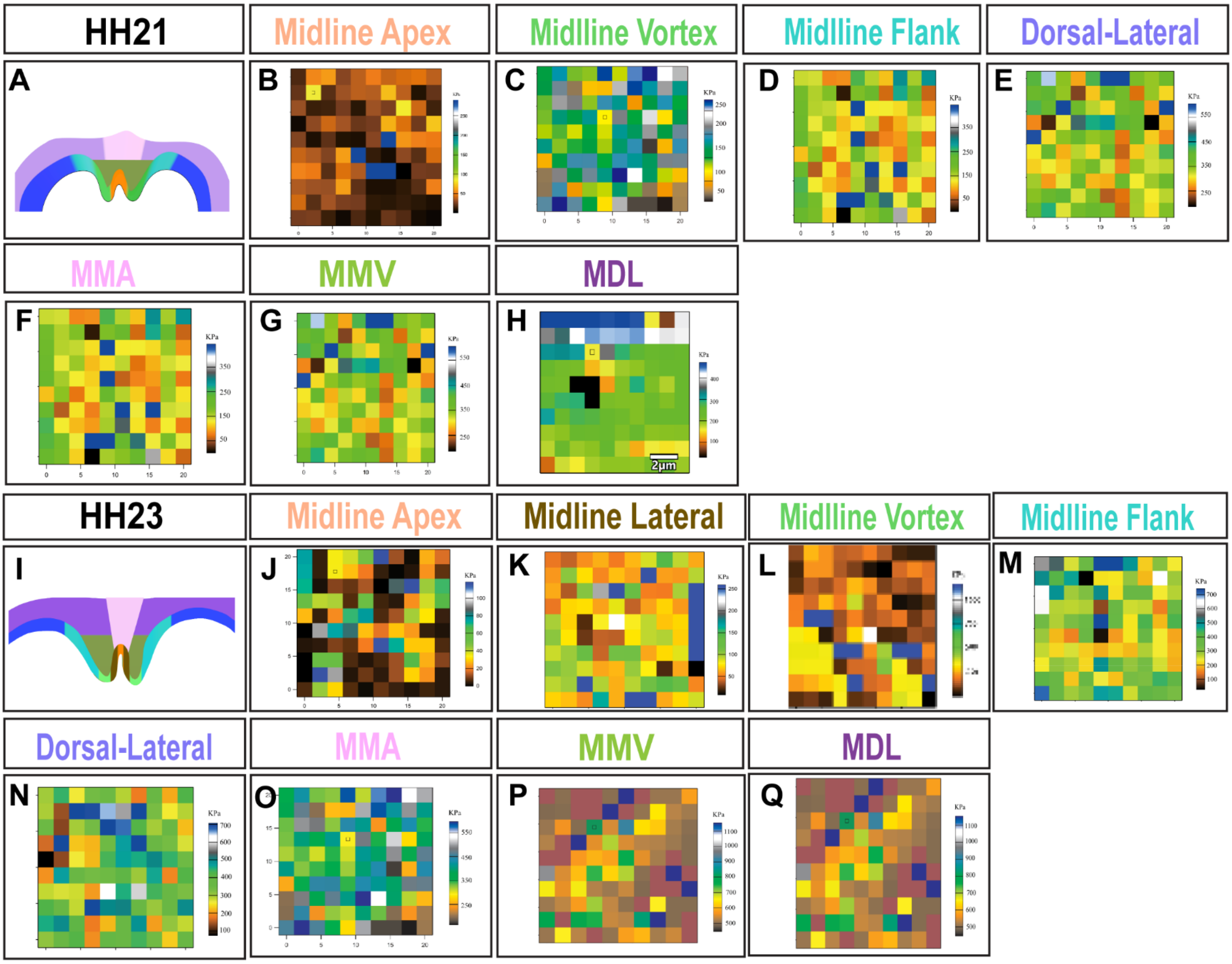


**Fig. S3. Heat maps for the measured Young's modulus in the chick forebrain at HH21 and HH23.**

(A) A schematic representation of a coronal slice from the middle posterior region of the chick dorsal forebrain at HH21 with distinct colours marking subdomains of the roof plate neuroepithelium and overlying mesenchyme.

(B,C,D,E) The representative heat maps for the subdomains of the neuroepithelium (Midline Apex, Midline Vortex, Midline Flank and Dorsal-Lateral) at HH21. For each region, a 20*20 μm² area was scanned to measure YM in triplicate.

(F,G,H) The representative heat maps for the subdomains of the mesenchyme (MMA, MMV and MDL) at HH21. For each region, a 20*20 μm² area was scanned to measure YM in triplicate.

(I) A schematic representation of a coronal slice from the middle posterior region of the chick dorsal forebrain at HH23 with distinct colours marking subdomains of the roof plate neuroepithelium and overlying mesenchyme.

(J,K,L,M,N) The representative heat maps for the subdomains of the neuroepithelium (Midline Apex, Midline Lateral, Midline Vortex, Midline Flank and Dorsal-Lateral) at HH21. For each region, a 20*20 $\mu m^2$ area was scanned to measure YM in triplicate.

(O,P,Q) ) The representative heat maps for the subdomains of the mesenchyme (MMA, MMV and MDL) at HH23. For each region, a 20*20 $\mu m^2$ area was scanned to measure YM in triplicate.

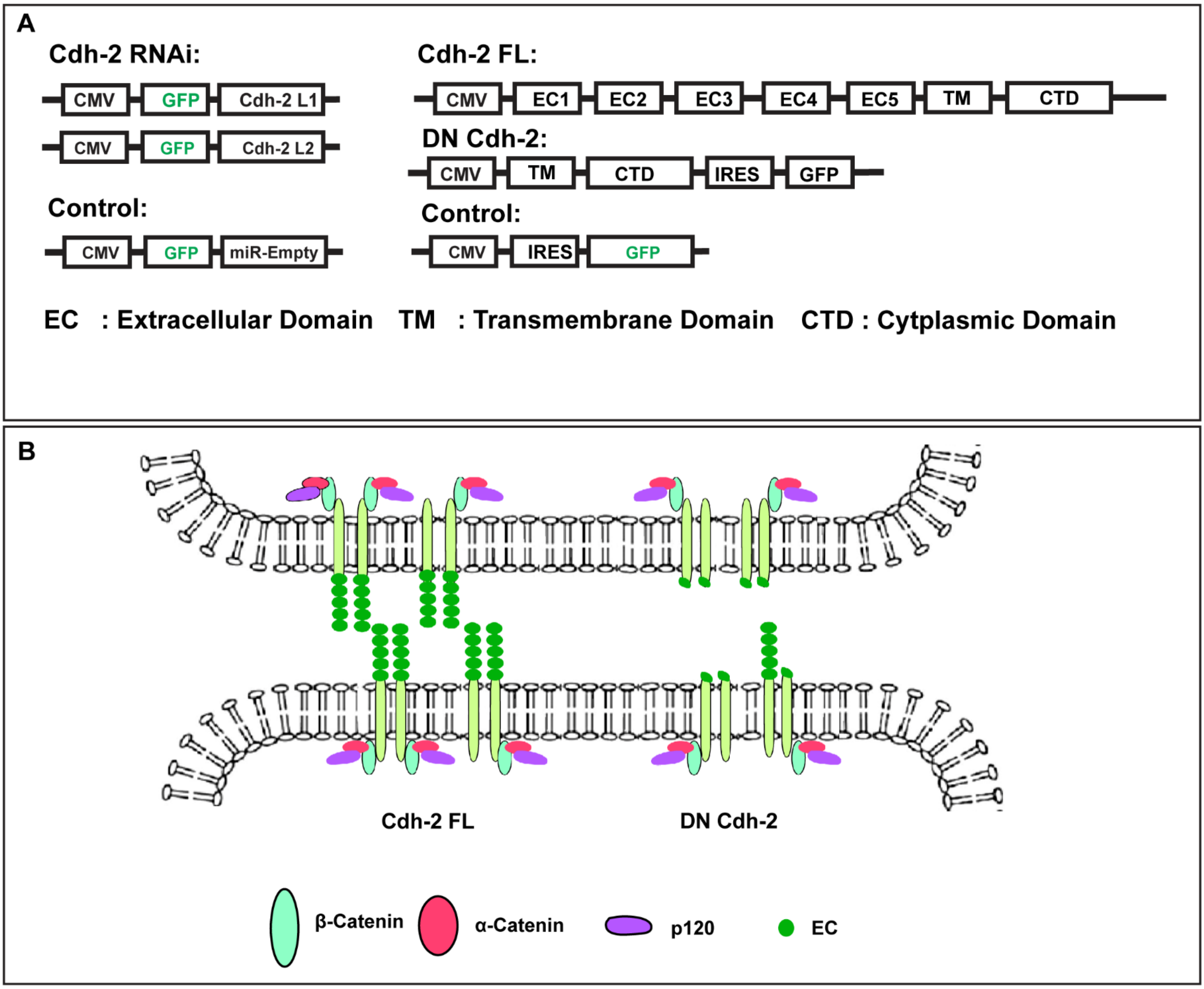


**Fig. S4. Design and schematic of Cdh-2 gain- and loss-of-function constructs.**

(A) Schematic of constructs used for loss of function and gain of function of Cdh-2.

(B) Strategy for creating loss of function of Cdh-2 through overexpression of dominant negative Cdh-2 (DN-Cdh-2). The schematic shows Cdh-2 with intact extracellular and intracellular domains, alongside a version of Cdh-2 that lacks the extracellular domain, resulting in a dominant negative variant (DN-Cdh-2) created by removing 5 tandem repeats from the extracellular domain.

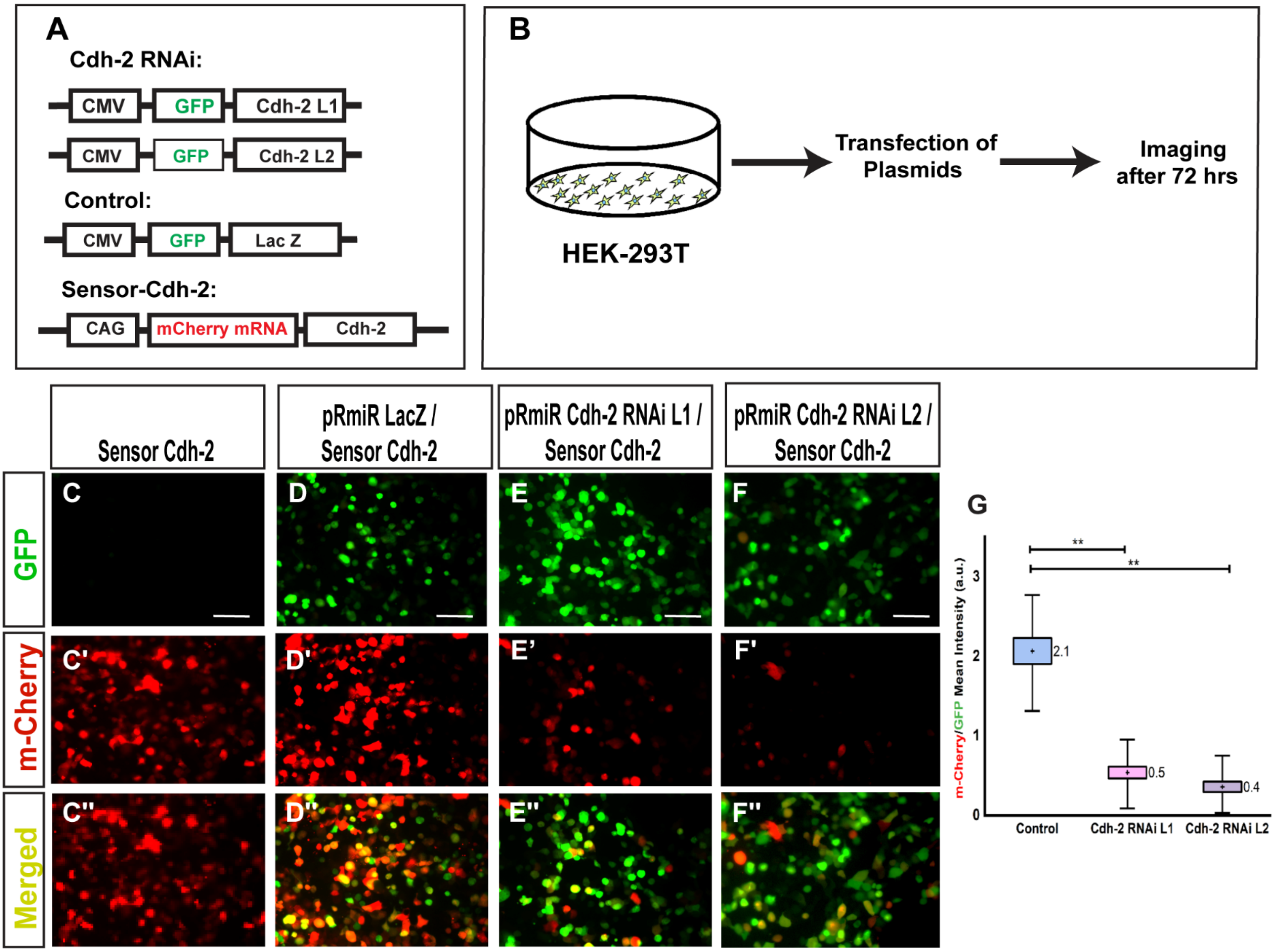


**Fig. S5. Sensor assay to determine the efficacy of Cdh-2 RNAi constructs.**

(A) Schematic representation of the constructs used to test the efficacy of RNAi oligos designed for knocking down Cdh-2 expression in HEK293T cells using the sensor assay.

(B) Schematic of the experimental design for the sensor assay.

(C, C' and C") HEK293T cells transfected with Sensor-Cdh-2 expressing mCherry alone. (C) GFP fluorescence (green), (C′) mCherry fluorescence (red) and (C″) Merged image for the HEK293T cells transfected with Sensor-Cdh-2 alone. Scale bar: 100μm

(D, D' and D") HEK293T cells co-transfected with Sensor-Cdh-2 and control RNAi-LacZ expressing GFP. (D) GFP fluorescence (green), (D′) mCherry fluorescence (red) and (D″) Merged images for the HEK293T cells co-transfected with Sensor-Cdh-2 and PRmiR-LacZ. Scale bar: 100μm

(E, E’ and E’’) HEK293T cells co-transfected with Sensor-Cdh-2 and PRmiR-Cdh-2 L1. (E) GFP fluorescence (green), (E′) mCherry fluorescence (red) and (E″) Merged images for the HEK293T cells transfected with Sensor-Cdh- 2 and PRmiR-Cdh-2 L1. Scale bar: 100µm

(F, F’ and F’’) HEK293T cells co-transfected with Sensor-Cdh-2 and PRmiR-Cdh-2 L2 expressing GFP. (F) GFP fluorescence (green), (F′) mCherry fluorescence (red) and (F″) Merged images for the HEK293T cells transfected with Sensor-Cdh-2 and PRmiR- Cdh-2-L2. Scale bar: 100µm

(G) Quantification of mean mCherry/GFP intensity for HEK293T cells transfected with only Sensor-Cdh-2, Sensor-Cdh-2 + PRmiR-Cdh-2 L1 and Sensor-Cdh-2 + PRmiR-Cdh-2-L2. Unpaired t-test using OriginPro software for determination of statistical significance. $p<0.01$ for all the comparisons. Error bars indicate mean±SEM. Scale bar: 100 µm for all experiments. Green fluorescence indicates cell where the RNAi construct effectively knocked down the sensor and yellow fluorescence indicates cells that have sensor expression (m-cherry) in the presence of RNAi (GFP) indicating a failure of knockdown. N=3 for all experiments.

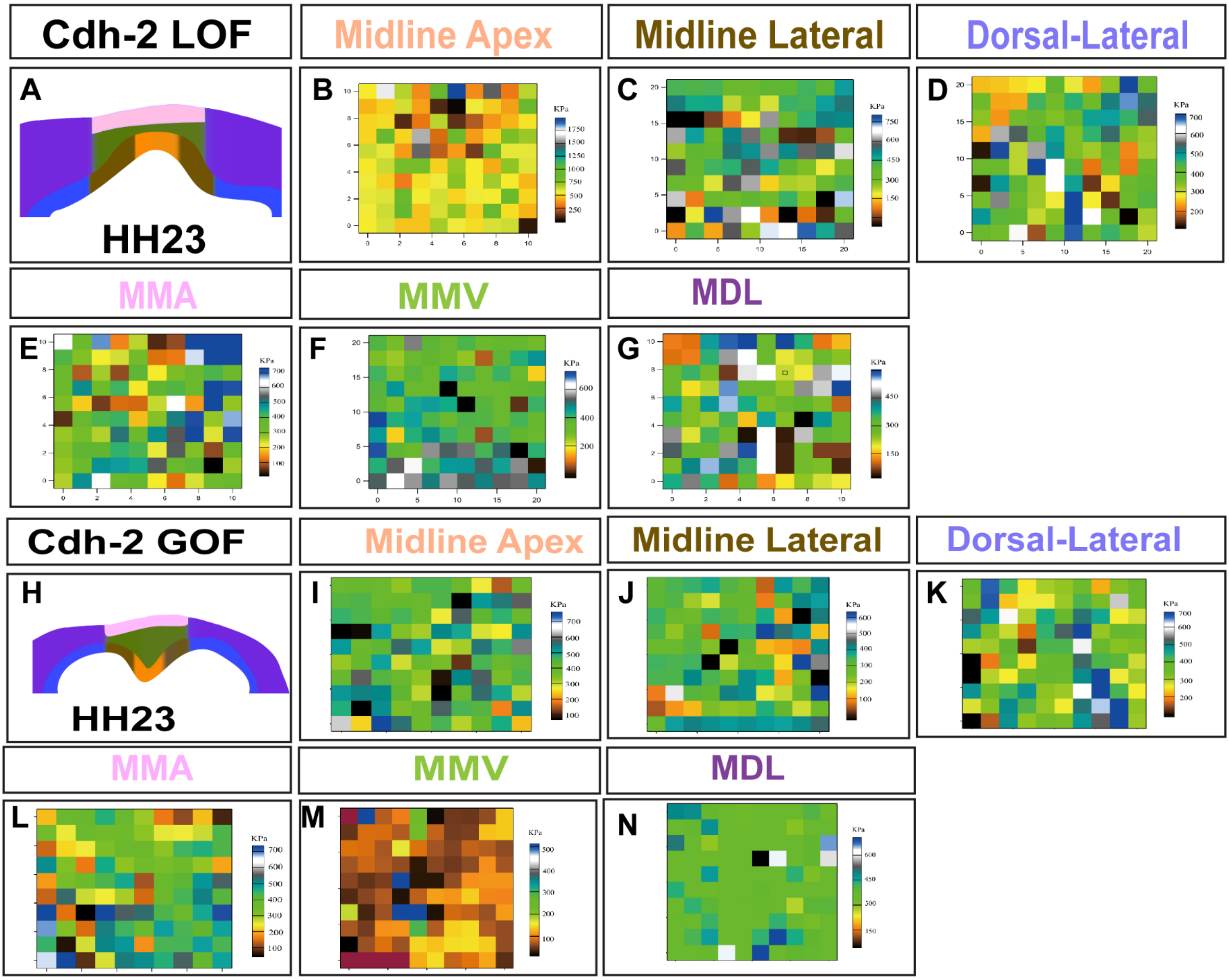


**Fig. S6. Heat maps for measured Young's modulus in the chick forebrain at HH23 following modulation of Cdh-2.**

(A) A schematic representation of a coronal slice from the middle posterior region of the chick dorsal forebrain upon LOF of Cdh-2 at HH23 with distinct colours marking subdomains of the roof plate neuroepithelium and overlying mesenchyme.

(B,C,D) The representative heat maps for the subdomains of the neuroepithelium (Midline Apex, Midline Lateral and Dorsal-Lateral) upon LOF of Cdh-2 at HH23. For each region, a 20*20 μm² area was scanned to measure YM in triplicate.

(E,F,G) The representative heat maps for the subdomains of the mesenchyme (MMA, MMV and MDL) upon LOF of Cdh-2 at HH23. For each region, a 20*20 μm² area was scanned to measure YM in triplicate.

(H) A schematic representation of a coronal slice from the middle posterior region of the chick dorsal forebrain upon GOF of Cdh-2 at HH23 with distinct colours marking subdomains of the roof plate neuroepithelium and overlying mesenchyme.

(I,J,K) The representative heat maps for the subdomains of the neuroepithelium (Midline Apex, Midline Lateral and Dorsal-Lateral) upon GOF of Cdh-2 at HH23. For each region, a 20*20 $\mu m^2$ area was scanned to measure YM in triplicate.

(L,M,N) The representative heat maps for the subdomains of the mesenchyme (MMA, MMV and MDL) upon GOF of Cdh-2 at HH23. For each region, a 20*20 $\mu m^2$ area was scanned to measure YM in triplicate.

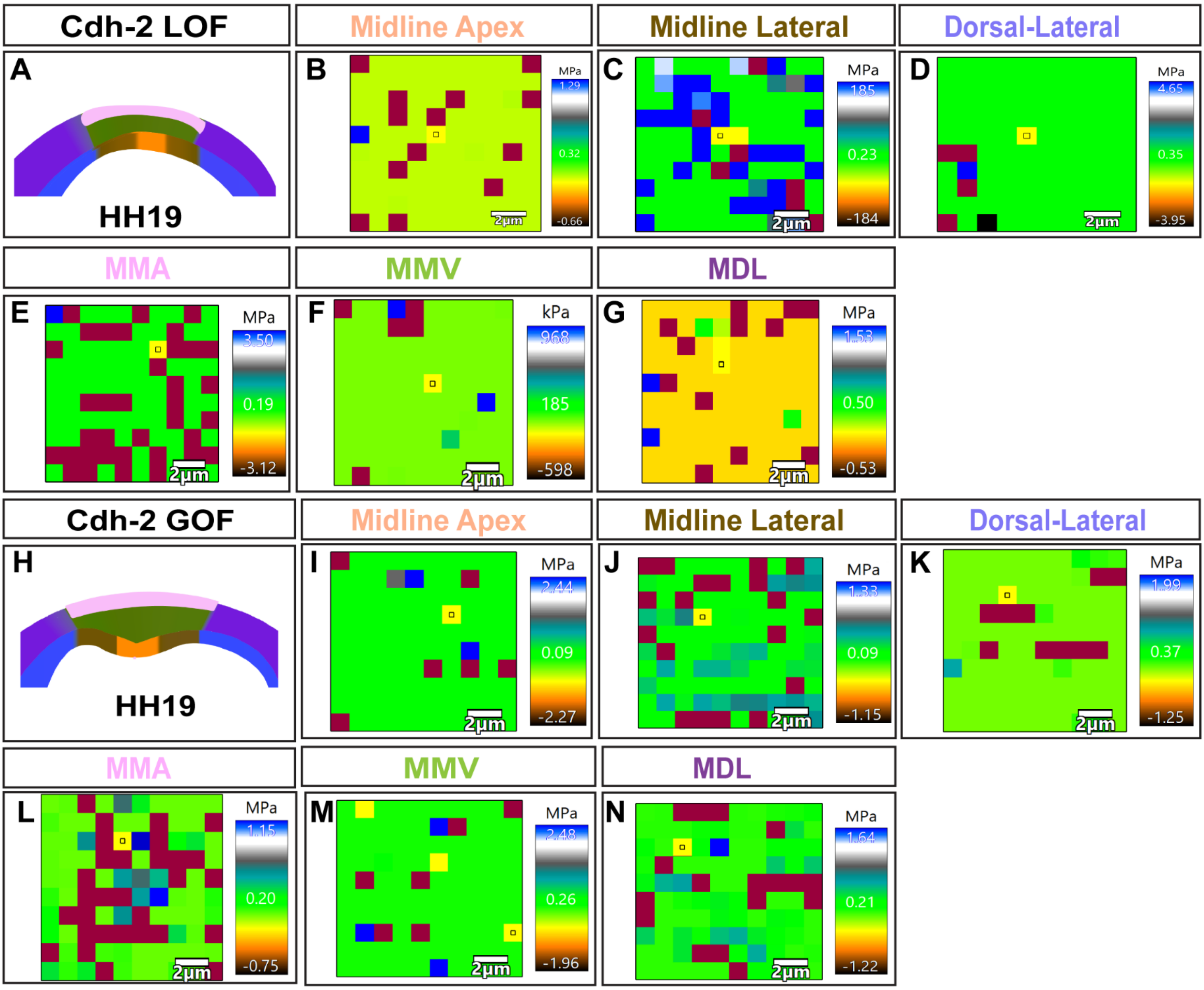


**Fig. S7. Heat maps for measured Young's modulus in the chick forebrain at HH19 following modulation of Cdh-2.**

(A) A schematic representation of a coronal slice from the middle posterior region of the chick dorsal forebrain upon LOF of Cdh-2 at HH19 with distinct colours marking subdomains of the roof plate neuroepithelium and overlying mesenchyme.

(B,C,D) The representative heat maps for the subdomains of the neuroepithelium (Midline Apex, Midline Lateral and Dorsal-Lateral) upon LOF of Cdh-2 at HH19. For each region, a 20*20 µm² area was scanned to measure YM in triplicate.

(E,F,G) The representative heat maps for the subdomains of the mesenchyme (MMA, MMV and MDL) upon LOF of Cdh-2 at HH19. For each region, a 20*20 µm² area was scanned to measure YM in triplicate.

(H) A schematic representation of a coronal slice from the middle posterior region of the chick dorsal forebrain upon GOF of Cdh-2 at HH19 with distinct colours marking subdomains of the roof plate neuroepithelium and overlying mesenchyme.

(I,J,K) The representative heat maps for the subdomains of the neuroepithelium (Midline Apex, Midline Lateral and Dorsal-Lateral) upon GOF of Cdh-2 at HH19. For each region, a 20*20 $\mu m^2$ area was scanned to measure YM in triplicate.

(L,M,N) The representative heat maps for the subdomains of the mesenchyme (MMA, MMV and MDL) upon GOF of Cdh-2 at HH19. For each region, a 20*20 $\mu m^2$ area was scanned to measure YM in triplicate.

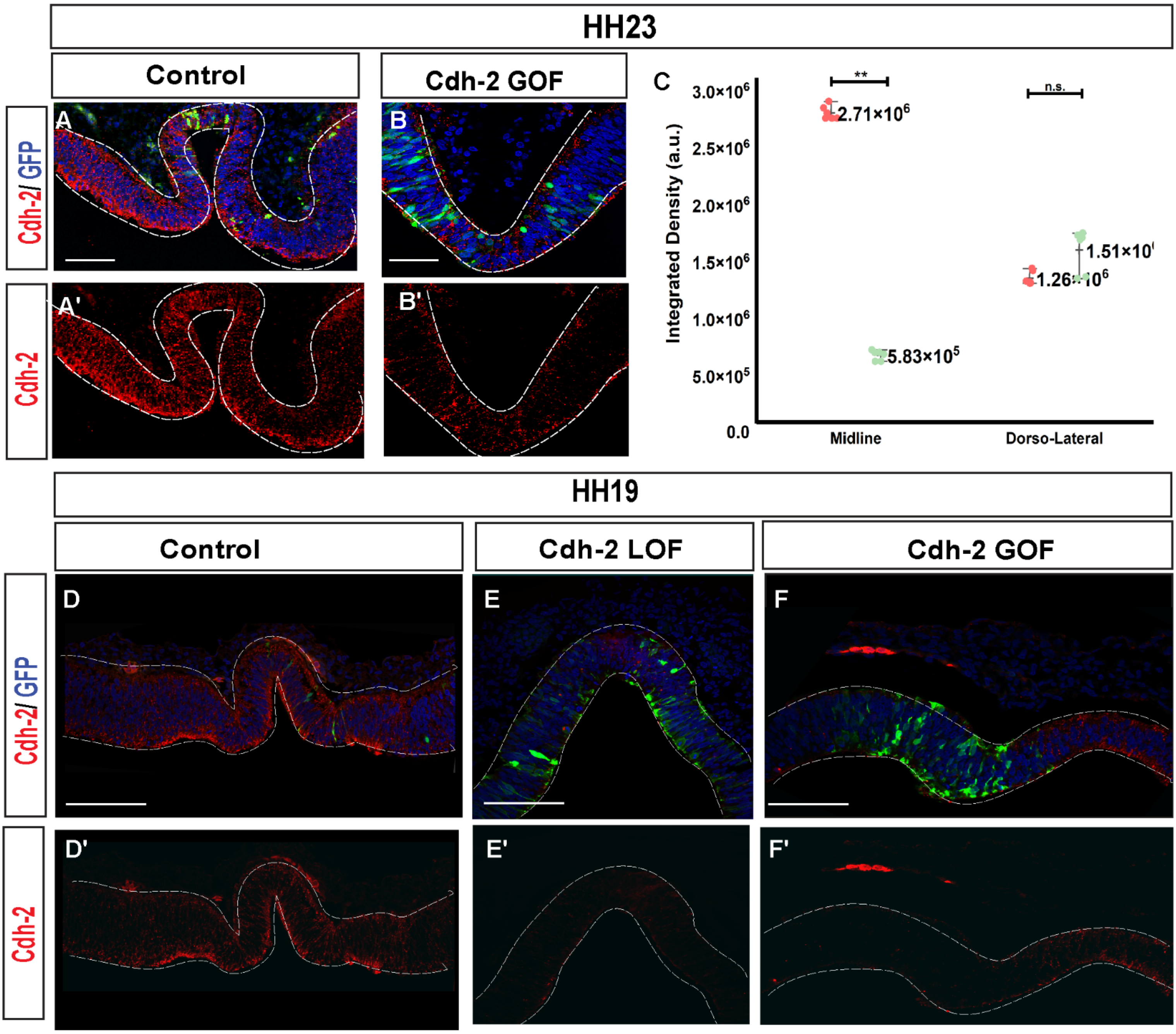


**Fig. S8. Impact of Cdh-2 manipulation on the levels of endogenous Cdh-2.**

Integration density of Cdh-2 was measured to quantify the amount of Cdh-2 protein and determine the effect of Cdh-2 overexpression (Cdh-2 GOF) in midline and dorsal lateral regions of neuroepithelium**.**

(A) Merged image of a section of chick forebrain electroporated with the control construct pCAG-GFP at HH23; DAPI (blue) marks nuclei, the green fluorescent signal demarcates the domain of electroporation, and red fluorescence indicates immunostaining of Cdh-2 protein. (A') Image of the same section as in (A) with red signal indicating Cdh-2 immunostaining. Scale bar: 100µm.

(B) The merged image shows a section of the chick forebrain co-electroporated with pCAG-GFP and Cdh-2 FL at HH23; DAPI (blue) marks the nuclei, the green fluorescent signal indicates the domain of electroporation, and the red fluorescence indicates immunostaining of Cdh-2 protein. (B') Image of the same section as in (B) with red signal indicating Cdh-2 immunostaining. Scale bar: 100µm.

3*3 boxes were marked in the midline and dorsal lateral regions in which the integrated density of Cdh-2 (red) was calculated.

(C) Dot plot comparing the integrated density of Cdh-2 (red) in the midline and dorsal lateral regions between the test and control groups. Error bars indicate mean±SEM, $p \leq 0.01$. N=6.

(D) Merged image of a section of chick forebrain electroporated with the control construct (pCAG-GFP) at HH19; DAPI (blue) marks nuclei, the green fluorescent signal demarcates the domain of electroporation, and red marks Cdh-2 immunostaining. (D') Image of the same section as in (D) with red signal indicating Cdh-2 immunostaining. Scale bar: 100µm.

(E) Merged image of a section of the chick forebrain electroporated with DN-Cdh-2 (LOF for Cdh-2) at HH19; DAPI (blue) marks nuclei, the green fluorescent signal demarcates the extent of electroporation, and red marks Cdh-2 immunostaining. (E') Image of the same section as in (E) with red signal denoting Cdh-2 immunostaining. Scale bar: 100µm.

(F) Merged image of a section of the chick forebrain co-electroporated with pCAG-GFP and Cdh-2 FL (GOF for Cdh-2) at HH19; DAPI (blue) marks nuclei, the green fluorescent signal demarcates the extent of electroporation, and red marks Cdh-2 immunostaining. (F') Image of the same section as in (F) with red signal denoting Cdh-2 immunostaining. Scale bar: 100µm.

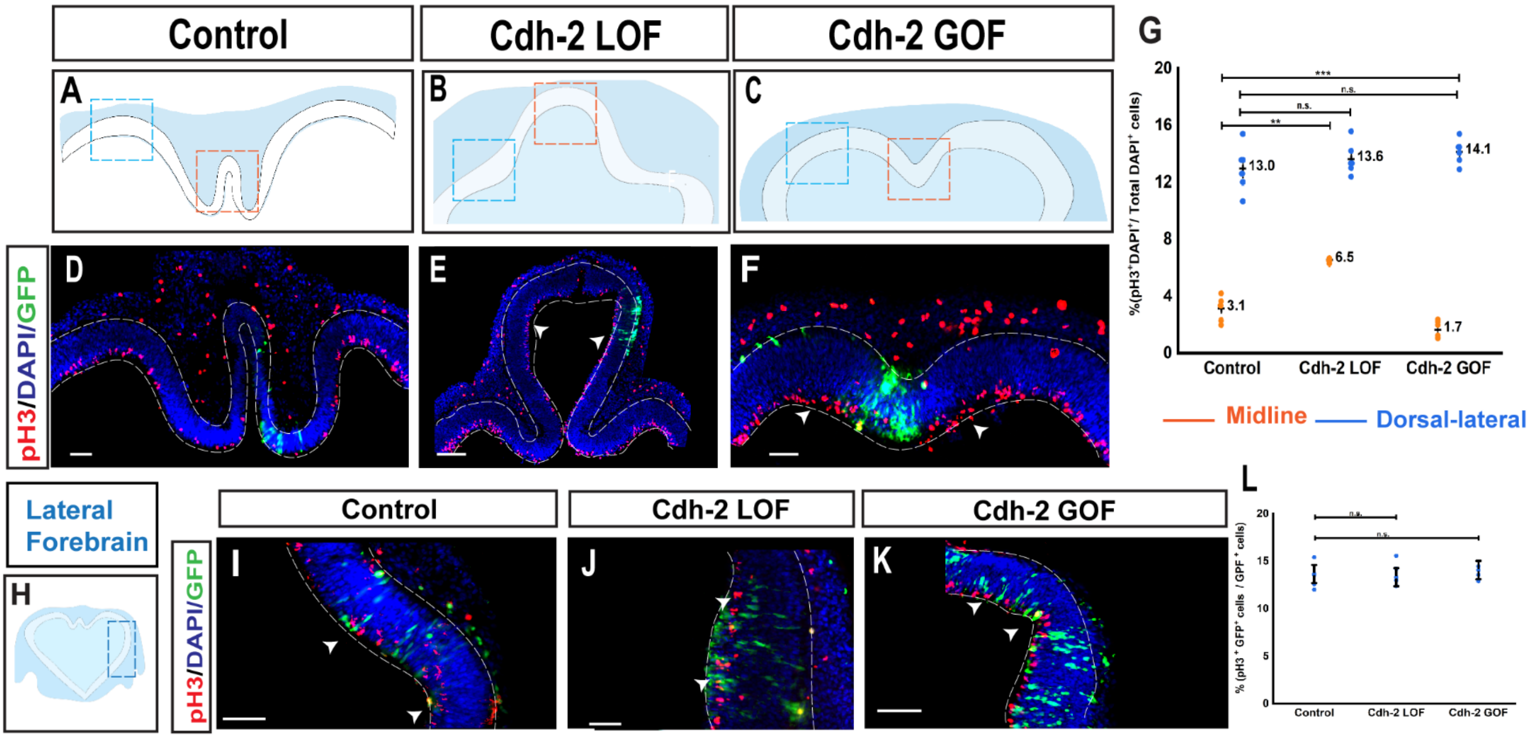


**Fig. S9. Effect of Cdh-2 modulation on the cell proliferation.**

(A, B, C) This schematic illustration depicts the region of the medial (orange boxed region) and dorsolateral (blue boxed region) neuroepithelium where PH3-positive cells were quantified to determine the effect of Cdh-2 perturbation on cell proliferation.

(D, E, F) A merged image shows a section of the chick forebrain roof plate that was electroporated with (D) control construct pCAG-GFP, E) DN-Cdh-2 construct and (F) Cdh-2 FL and pCAG-GFP. The nuclei of neuroepithelial cells are identified by DAPI staining (blue), and the domain of electroporation is indicated by green fluorescence. Red fluorescence indicates PH3-positive proliferating cells. Scale bar: 100µm.

(G) Quantification of percentage of DAPI-positive cells that are PH3 positive, statistical significance determined using an unpaired t-test, Origin 2024b. (N=5), $p \leq 0.001$.

(H) Schematic of the section of the chick forebrain at HH23 with the blue dashed box indicating the position of the lateral region of the neuroepithelium used to quantify PH3 and DAPI.

(I, J, K) Merged image of a section of the chick forebrain roof plate electroporated with (I) the control construct IRES-GFP, (J) the DN-Cdh-2 construct and (K) the Cdh-2 FL and pCAG-GFP constructs where DAPI staining (blue) marks the nuclei of neuroepithelial cells, green fluorescence demarcates the domain of electroporation and  Red fluorescence marks the PH3-positive proliferating cells. Scale bar: 100µm.

(L) Box plot showing quantification of the percentage of double positive (PH3+ and GFP+) cells by total GFP+ positive cells, unpaired t-test using Origin 2024b software was carried out for statistical significance. (N=4), $p \leq 0.001$.

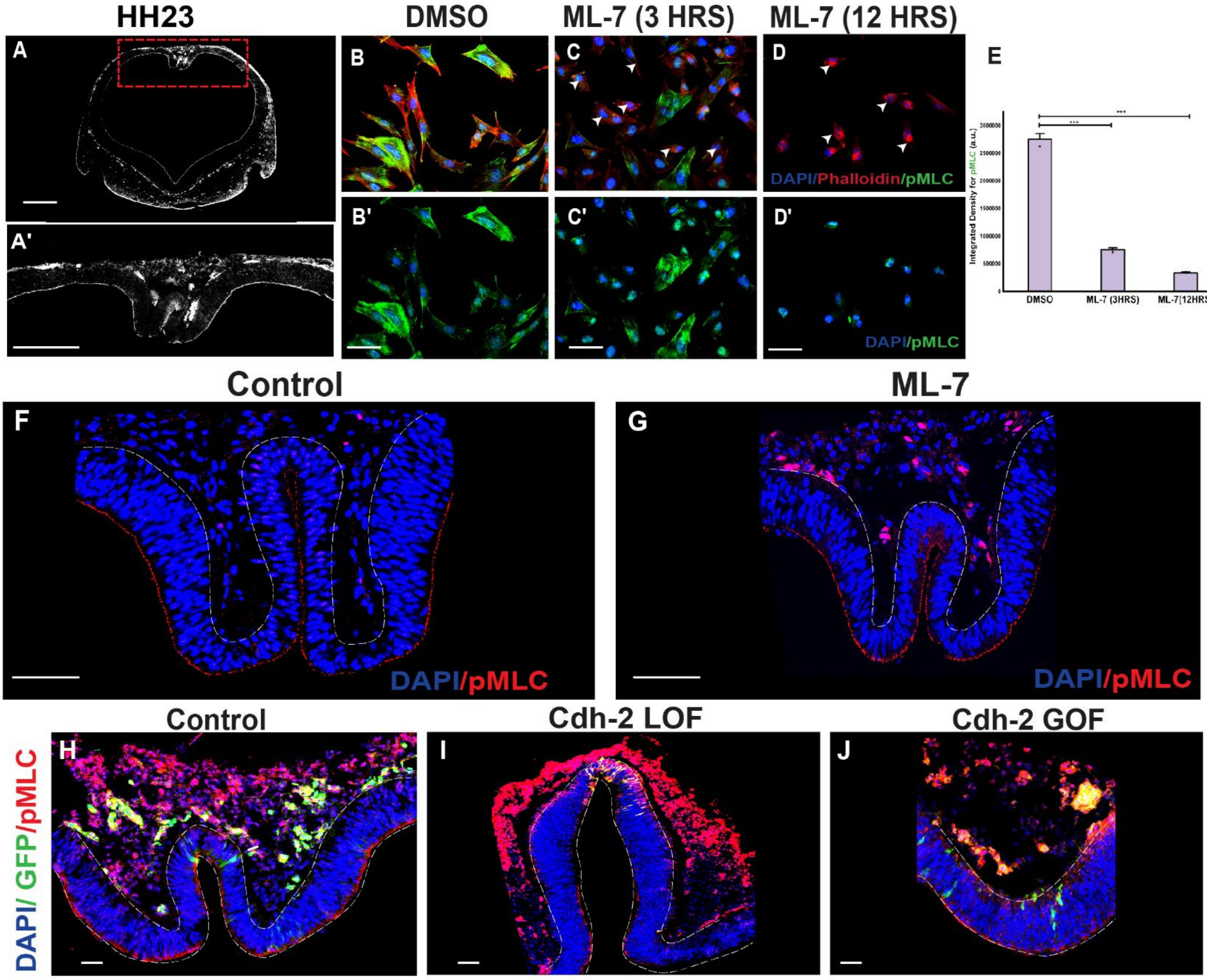


**Fig. S10. Effect of pMLC modulation on forebrain roof plate and effect of Cdh-2 modulation on pMLC.**

(A) & (A') Immunohistochemical staining of pMLC (Grey) on the section of the forebrain at HH23 (embryonic day 3.5). To evaluate the efficacy of ML-7, we assessed the effect of the inhibitor on DF1 cells. For this (B) DF1 cells treated with DMSO (50μM); DAPI (blue) marks nuclei, the green fluorescent signal demarcates pMLC, and red fluorescence indicates phalloidin marking F-actin. Scale bar: 100μm. (B') Merged images for the DF1 treated with DMSO. DAPI (blue) marks nuclei, the green fluorescent signal demarcates pMLC. (C) DF1 cells treated with ML-7 (50μM) for 3 Hrs ; DAPI (blue) marks nuclei, the green, fluorescent signal demarcates pMLC, and red fluorescence indicates phalloidin marking f-actin. (C') Merged images for the DF1 treated with ML-7 (50μM) for 3 Hrs. DAPI (blue) marks nuclei, the green, fluorescent signal demarcates pMLC. (D) DF1 cells treated with ML-7 (50μM) for 12 Hrs ; DAPI (blue) marks nuclei, the green fluorescent signal demarcates pMLC, and red fluorescence indicates phalloidin marking F-actin. (D') Merged images for the DF1 treated with ML-7 (50μM) for 3 Hrs. DAPI (blue) marks nuclei, the green, fluorescent signal demarcates pMLC. Scale Bar -100 μm. (E) Quantification of mean integrated density for pMLC in DF1 cells treated with only DMSO, ML-7 for 3Hrs and 12 Hrs. Unpaired t-test using OriginPro software for determination of statistical significance. $P<0.001$ for all the comparisons. Error bars indicate mean±SEM. Scale bar: 100 µm for all experiments. N=3 for all experiments.(F) Merged image of a section of the chick forebrain treated with DMSO (50 µM) at HH23; DAPI (blue) marks nuclei and the red marks pMLC immunostaining. (G) Merged image of a section of the chick forebrain treated with ML-7 (50 µM) at HH23; DAPI (blue) marks nuclei and the red marks pMLC immunostaining. (H) Merged image of a section of chick forebrain electroporated with the control construct (pCAG-GFP) at HH23; DAPI (blue) marks nuclei, the green fluorescent signal demarcates the domain of electroporation, and red marks pMLC immunostaining. (I) Merged image of a section of the chick forebrain electroporated with DN-Cdh-2 (LOF for Cdh-2) at HH23; DAPI (blue) marks nuclei, the green fluorescent signal demarcates the extent of electroporation, and red marks pMLC immunostaining. (J) Merged image of a section of the chick forebrain co-electroporated with pCAG-GFP and Cdh-2 FL (GOF for Cdh-2) at HH23; DAPI (blue) marks nuclei, the green fluorescent signal demarcates the extent of electroporation, and red marks pMLC immunostaining. Scale bar: 100µm.

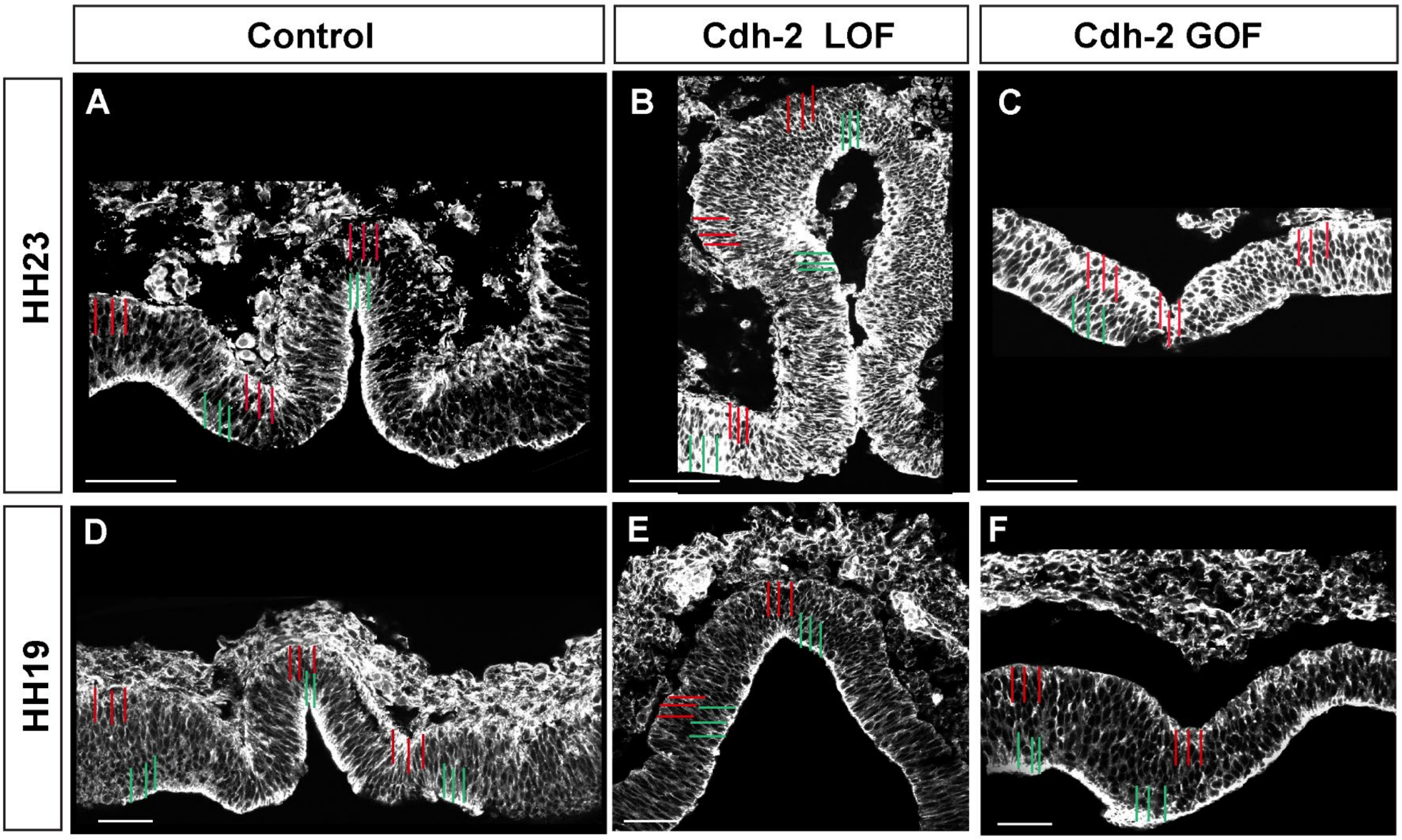


**Fig. S11. Effect of Cdh-2 modulation on cortical F-actin distribution in neuroepithelial cells.**

(A, B, C) Fluorescence intensity of phalloidin was measured up to 80 a.u into the cells from both the apical and basal surfaces (orange and green lines) for sections of the forebrain roof plate at HH23 electroporated with (A) the control construct, (B) the LOF for Cdh-2 construct and (C) the GOF for Cdh-2 construct at HH23. The green lines mark the neuroepithelium on the apical surface, while the orange lines mark the neuroepithelium on the basal surface. The intensity was measured at the midline apex (MA), midline vortex (MV), and dorsal lateral (DL) regions of the neuroepithelium. Scale bar: 100µm.

(D, E, F) Fluorescence intensity of phalloidin was measured up to 80 a.u into the cells from both the apical and basal surfaces (orange and green lines) for sections of the forebrain roof plate at HH19 electroporated with (D) the control construct, (E) the LOF for Cdh-2 construct and (F) the GOF for Cdh-2 construct at HH23. The green lines mark the neuroepithelium on the apical surface, while the orange lines mark the neuroepithelium on the basal surface. The intensity was measured at the midline apex (MA), midline vortex (MV), and dorsal lateral (DL) regions of the neuroepithelium. Scale bar: 100µm.
